# A Wearable Optical Microfibrous Biomaterial with Encapsulated Nanosensors Enables Wireless Monitoring of Oxidative Stress

**DOI:** 10.1101/2020.07.24.220194

**Authors:** Mohammad Moein Safaee, Mitchell Gravely, Daniel Roxbury

**Affiliations:** Department of Chemical Engineering, University of Rhode Island, 2 East Alumni Avenue, Kingston, Rhode Island 02881, United States

**Keywords:** wearable biosensors, wound healing, wound monitoring, single-walled carbon nanotubes, biomaterials, oxidative stress, infection

## Abstract

In an effort to facilitate personalized medical approaches, the continuous and noninvasive monitoring of biochemical information using wearable technologies can enable a detailed understanding of an individual’s physiology. Reactive oxygen species (ROS) are a class of oxygen-containing free radicals which function in a wide range of biological processes. In wound healing applications, the continuous monitoring of ROS through a wearable diagnostics platform is essential for the prevention of chronicity and pathogenic infection. Here, a versatile one-step procedure is utilized to fabricate optical core-shell microfibrous textiles incorporating single-walled carbon nanotubes (SWCNTs) for the real-time optical monitoring of hydrogen peroxide concentrations in wounds. The environmentally sensitive and non-photobleachable fluorescence of SWCNTs enables continuous analyte monitoring without a decay in signal over time. The existence of multiple chiralities of SWCNTs emitting near-infrared fluorescence with narrow bandwidths allows a ratiometric signal readout invariant to the excitation source distance and exposure time. The individual fibers encapsulate the SWCNT nanosensors for at least 21 days without apparent loss in structural integrity. Moreover, the microfibrous textiles can be utilized to spatially resolve peroxide concentrations on a wound surface using a camera and can be integrated into commercial wound bandages without being altered or losing their optical properties.

## 1. Introduction

Reactive oxygen species (ROS) are continuously generated and consumed in all eukaryotic and prokaryotic cells as a consequence of aerobic life.^[1, 2]^ In this biological context, ROS primarily function to preferentially react with specific atoms of biomolecules involved in a wide range of physiological processes.^[1]^ ROS play a crucial role in biological signaling including the inhibition or activation of proteins, subsequent promotion or suppression of inflammation, immunity, and carcinogenesis.^[1, 3–6]^ Oxidative stress can occur if the ROS-induced signal is too strong, if it persists for too long, or if it occurs at the wrong time or place.^[1, 3, 5, 7]^ As a key example, wound healing is one of the most dynamic biological processes involving ROS-linked cellular signaling throughout the entire mechanism.^[8–11]^ Additionally, basal concentrations of ROS aid in the fight against invading microorganisms into open wounds.^[4, 11, 12]^ The excessive and uncontrolled production of ROS contributes to the sustaining and deregulation of inflammation processes, which play a central role in the pathogenesis of chronic non-healing wounds.^[13–17]^ Physiologically, hydrogen peroxide (referred herein as peroxide) and superoxide function as intracellular ROS messengers stimulating key phases of wound healing including cell recruitment, production of cytokines, and angiogenesis.^[16, 18, 19]^ Of note, peroxide acts as the principal secondary messenger in wound healing and is present at low concentrations (100–250 μM) in normal wounds.^[8, 11, 19, 20^ Increased peroxide concentration is a biomarker for inflammation and chronicity in which biofilm-forming pathogens can grow significantly faster than acute wounds.^[15, 17, 20]^ Interestingly, strictly controlling the ROS levels through antioxidants has recently been shown to improve inflammatory skin conditions and wound healing process in diabetic and hypoxic environments.^[21]^

Due to the short half-lives of ROS, the direct detection and quantification of their concentrations are often difficult in the laboratory and in patients.^[20]^ Although multiple classes of sensors and spectrophotometric assays have been developed to monitor various types of ROS, the current methods are limited in their capabilities. Spectrophotometric methods,^[22]^ such as total antioxidant capacity assays (TAC),^[23]^ and gel electrophoresis^[14]^ have been utilized to indirectly determine the oxidation products of lipids, proteins and DNA, but these are not capable of real-time monitoring in the wound site.^[20]^ Various label-free electrochemical biosensors have been developed to accurately quantify the ROS concentrations by immediately converting the chemical information to an electrical signal.^[19, 24, 25]^ The main drawback of electrochemical techniques is the requirement to incorporate electrodes into different biomaterials and wireless platforms. Moreover, the need to utilize an electrochemical signal transducer restricts the application of the current sensors on wounds in different organs of the body. Fluorescent nanoparticles^[26–28]^ and genetically encoded fluorescent molecules^[8, 29–31]^ highly selective for peroxide have been created to study the redox events in mice, zebrafish, and cells, but these assays cannot be utilized for real-time monitoring in clinical applications. Therefore, developing a point-of-care diagnostic technology for the real-time monitoring of ROS concentrations in wound sites is essential to prevent chronicity and infection, and to deliver accurate amounts of antioxidants and antibiotics to the wounds.

Single-walled carbon nanotubes (SWCNTs) with engineered wrappings have recently been developed and utilized in various disparate fields ranging from additives that strengthen material composites^[32, 33]^ to biomedical applications including near-infrared (NIR) optical biosensing,^[34–36]^ and biological imaging.^[37, 38]^ The electronic band gap energies of SWCNTs are dependent on their chiral identity, denoted by integers (n,m), and vary based on diameter and rollup angle, resulting in various semiconducting species which exhibit a distinct narrow-bandwidth photoluminescence in the second NIR window.^[39]^ The SWCNT photoluminescence responds to their local environment,^[40]^ resulting in shifts in emission wavelengths^[40–43]^ and/or variations in intensity.^[44, 45]^ Certain amphiphilic polymers such as short single-stranded deoxyribonucleic acids (ssDNA),^[41]^ Phospholipid-Polyethylene glycol (PL-PEG),^[46]^ and synthetic polymers^[34]^ have all been shown to effectively solubilize SWCNTs, imparting enhanced biocompatibility and desirable fluorescence properties. The resultant hybrid nanomaterials have been optimized for the detection of a wide range of analytes *in vivo* and *in vitro* including neurotransmitters,^[45]^ lipids,^[35]^ and proteins.^[43]^ SWCNT-based optical nanosensors have also recently been developed for real-time spatial and temporal monitoring of ROS in various plant species as a biomarker for plant health.^[44, 47]^ Moreover, ratiometric SWCNT-based optical sensors have enabled the real-time monitoring of ROS in plants, allowing an absolute calibration independent of overall intensity.^[48]^ The current ratiometric sensing approaches based on SWCNTs require separation of at least two highly pure single chiralities, wrapped in two different polymers, where one polymer-chirality pair is sensitive to the local environment and the other pair does not spectrally respond to the variations in the local environment and acts as a reference.^[48]^

Although the ssDNA- and polymer-wrapped SWCNT nanosensors have attracted significant interest in the past decade for biosensing applications *in vivo* and *in vitro*, the integration of these biosensors into other bulk biomaterial platforms has been a challenge as their NIR fluorescence is remarkably sensitive to the chemistry of their local environment and can be suppressed by other components in the biomaterial preparation processes.^[34, 40]^ Moreover, due to the hydrophilicity of these nanosensors, it is unfavorable to engage them in any process involving organic solvents as they form bulk aggregates in hydrophobic environments.

With revolutionary advances in nanotechnology and biomaterials in recent years, an extensive range of smart wound care biomaterials have been developed that enable localized delivery of drugs on the wound site^[49, 50]^ and real-time monitoring of the wound microenvironment.^[51, 52]^ Electrospun microfibers are one of the novel classes of wound dressings as they mimic the chemical and mechanical environment of the 3D extracellular matrix.^[53, 54]^ Microfiber-based wound dressings have been designed to enhance cell migration,^[55]^ prevent inflammation and infection,^[56]^ and inhibit scar formation on wounds.^[54]^ Herein, we utilized a one-step co-axial electrospinning process to fabricate wearable microfibrous textiles incorporating peroxide-sensing SWCNTs. The electrospun fibers feature a core-shell morphology in which the SWCNTs are encapsulated inside of a polymer shell that is soluble in an organic solvent. We chose polycaprolactone (PCL) as the shell material as it is an FDA-approved polymer which has been extensively studied for tissue engineering and wound healing applications.^[57–59]^ Our wearable optical platform was able to wirelessly and reversibly detect peroxide in a physiologically-relevant range for wounds (1-250 μM). The ratiometric characteristic of the NIR fluorescence sensor facilitates *in vivo* and clinical applications as it transduces an absolute signal that is not dependent on excitation source distance nor exposure time. Utilizing confocal Raman microscopy, we found that the SWCNT nanosensors stay encapsulated within the individual fibers for up to at least 21 days, indicating that the long-term identity of the nanosensing platform is maintained. We also indicate the potential of our optical textiles for spatially resolving the peroxide concentration on the surface of a wound using hyperspectral fluorescence microscopy. Finally, we attached our microfibrous platform to a conventional wound bandage and demonstrated the feasibility of *in situ* measurements of peroxide in wounds.

## 2. Results and Discussion

### 2.1. Preparation and Characterization of Optical Microfibers

We prepared aqueously-dispersed ssDNA-SWCNT nanosensors by probe-tip sonicating HiPco SWCNTs in the presence of single-stranded (GT)_15_ DNA (Figure 1a). The (GT)_15_ sequence was selected as SWCNT-based nanosensors with this sequence of DNA have been utilized in live cells and plants for the real-time and selective monitoring of peroxide, in contrast to other signaling molecules including nitric oxide (NO), super oxide ([O_2_]^•–^), singlet oxygen (^1^O_2_) and hydroxyl radical ([OH]^•^).^[27, 44, 48]^ Following sonication, the sample was ultracentrifuged to remove bundles of undispersed SWCNTs as well as residual catalyst particles to produce an ink-like solution with strong NIR absorbance and fluorescence spectra (Figure S1). With colloidally stable nanosensors, we employed a core-shell electrospinning procedure to encapsulate the hydrophilic ssDNA-SWCNTs along with poly(ethylene oxide) (PEO) into the polymer polycaprolactone (PCL), that is soluble in an organic solvent (Figure 1b). Briefly, the shell and core are extruded from a two-compartment spinneret, and once injected, form a core-shell pendant droplet as the result of surface tension.^[54]^ A high voltage is applied to the droplet that produces a two-compartment Taylor cone as well as a constant electric field between the spinneret and a metal grounded collector.^[54]^ The electrical force significantly elongates the two components of the Taylor cone until they turn into microfibers.^[54]^ After rapid solvent evaporation, the immiscibility of the core and shell causes complete encapsulation of the core within the shell.^[60]^ In this process, the hydrophilic ssDNA-SWCNT nanosensors are protected against a prolonged interaction with an organic solvent. Additionally, the intrinsic NIR fluorescence of the nanosensors is maintained as the process does not introduce any other chemicals such as crosslinking agents. There are a number of physical parameters involved in the electrospinning process which control the reproducibility and homogeneity of the final samples. We have optimized the flow rates of the polymers, rotation rate of the collector, and the distance between the needle and collector to achieve a stable electrospinning jet (data not shown).

**Figure 1.**
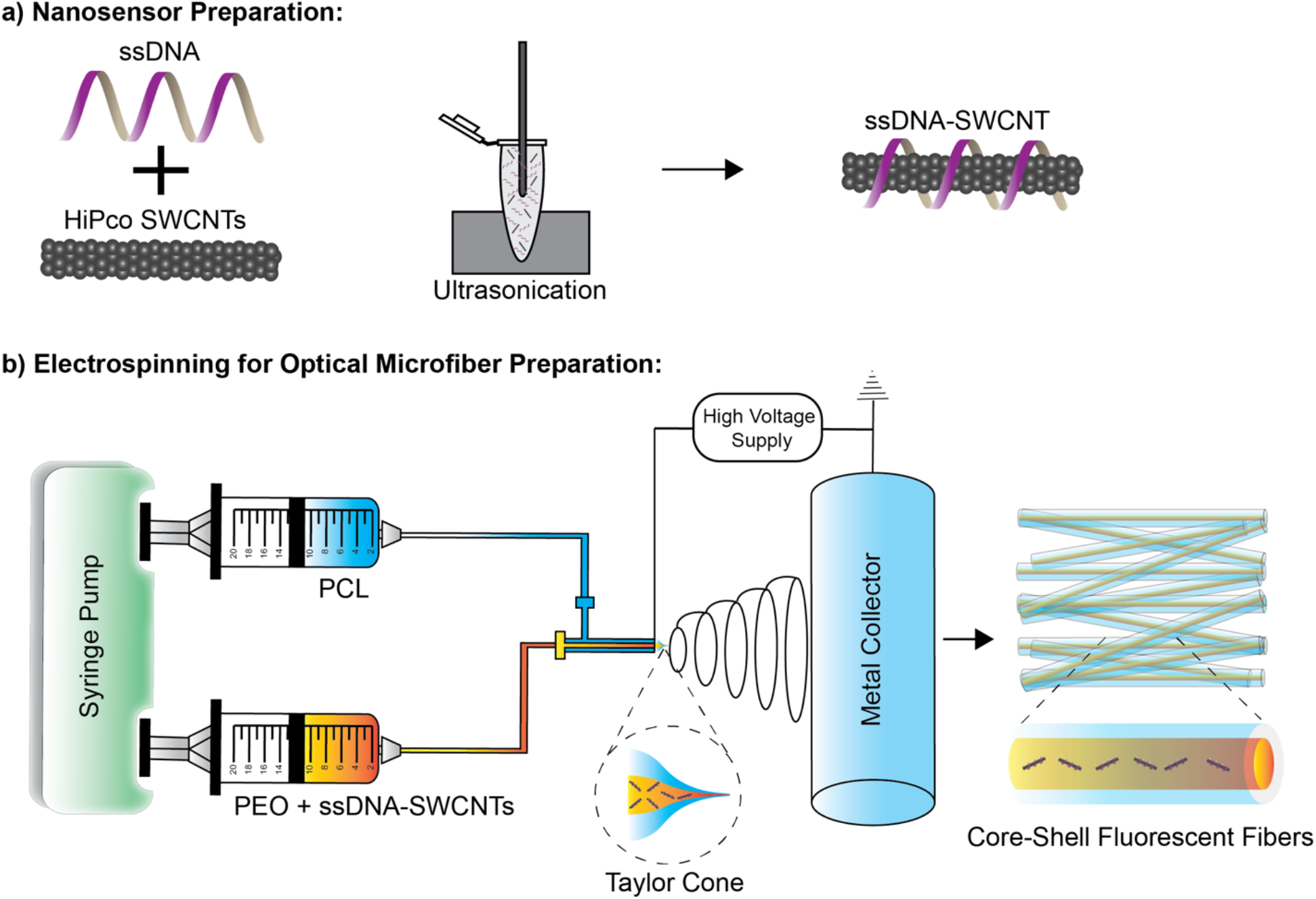
Fabrication of optical microfibers. (a) Nanosensor preparation by probe-tip sonicating SWCNTs in the presence of ssDNA followed by ultracentrifugation of the resultant dispersion. (b) Core-shell electrospinning setup for fabrication of the optical microfibrous textiles. The two syringes containing the core and shell polymer solutions are connected to the inlets of a custom core-shell needle. Once the polymer solutions are injected out, a core-shell pendant droplet is formed. A high-voltage supply is connected to the tip of the needle and electrifies the droplet, forms a Taylor cone, and eventually elongates the cone until microfibers are created. The resultant fibers are collected onto a rotating metal grounded collector.

To optimize the morphology of the fibers and aggregation state of the nanosensors, we have tuned the applied voltage during the electrospinning process. Figure 2 demonstrates NIR broadband fluorescence images (900-1400 nm) and scanning electron microscopy (SEM) images of fibers produced with three different voltages. The applied voltage of 12 kV did not provide a high enough rate of elongation, and as a result, SWCNT aggregates appeared in the NIR fluorescence images (Figure 2a and S2). When the applied voltage was 16 kV, occasional aggregates again emerged along the fibers (Figure 2c and S4), presumably due to incomplete formation of the Taylor cone. An applied voltage of 14 kV produced a homogenous fiber morphology with no significant spatially localized aggregations (Figure 2b and S3). The SEM images of the fibers produced with the three voltages revealed two subsets of fibers with diameters of more than ~1 μm or less than ~100 nm (Figure 2d-f, S5). By visually comparing the SEM and NIR fluorescence images, we observe that the diameter range of the NIR optical fibers is identical to the micron-size fibers in the SEM images (Figure S2-5). Thus, although we have produced a matrix of micro- and nanofibers, we acknowledge the fact that the optically-active fibers have sizes of more than 1 μm.

**Figure 2.**
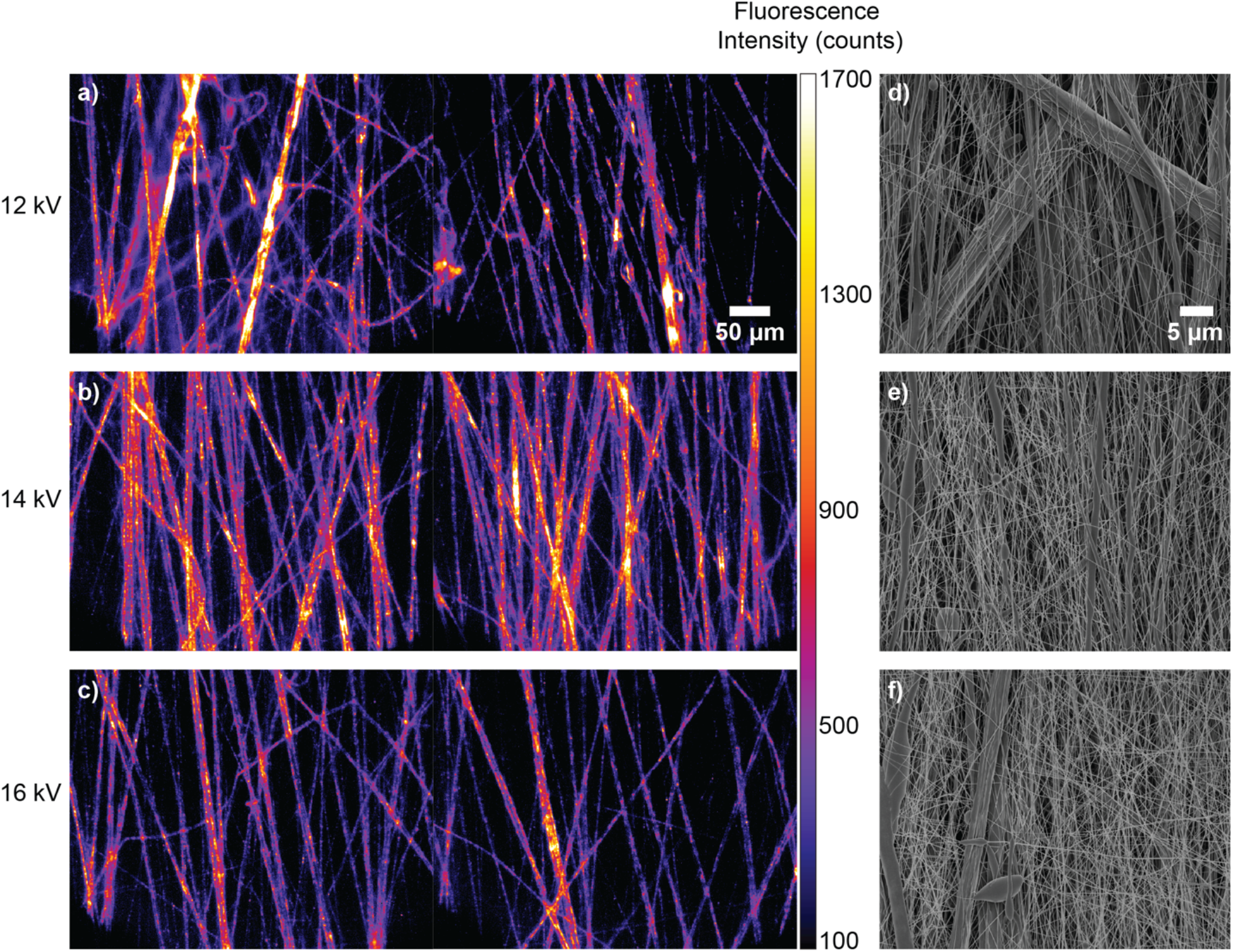
NIR broadband fluorescence images of microfibers produced with the applied voltages of (a) 12 kV, (b) 14 kV, and (c) 16 kV. The microfibers were illuminated with a 730 nm laser and images were acquired with a 2D InGaAs array detector in the wavelength range 900-1400 nm. SEM images of the fibers fabricated with applied voltages of (d) 12 kV, (e) 14 kV, and (f) 16 kV.

We employed confocal Raman microscopy to confirm the core-shell morphology and assess the complete encapsulation of nanosensors within the individual fibers. Confocal Raman microscopy is a powerful technique used to analyze multicomponent material samples, in which the unique Raman spectrum of each component can be identified and spatially resolved. SWCNTs exhibit distinct Raman signatures such as the G-band (1589 cm^−1^), which proportionately scales with increasing amounts of graphitic carbon (i.e. SWCNT concentration),^[61, 62]^ and the radial breathing mode (150 - 350 cm^−1^), which can identify the chiral composition of a SWCNT mixture.^[63, 64]^ Figure 3a displays the brightfield and G-band intensity overlay of a single as-produced fiber. A k-means clustering analysis was applied to the entire dataset, in which the spectrum from each pixel was partitioned into one of 4 clusters (*k* = 4) based on the location and intensity of individual peaks, creating 4 average spectra which best represented all regions, including the background, from the Raman area scan (Figure 3b and c).^[65]^ Based on the clustering analysis, each individual fiber was categorized to three areas: core, intermediate, and shell (background constituted the fourth cluster). The average Raman spectrum of the core area corroborated previous reports of SWCNT Raman spectra (Figure 3c).^[32, 41]^ The average Raman spectrum of the shell area predominantly matched with the spectrum from the PCL polymer (Figure S6a) with two additional peaks at ~1589 and 240 cm^−1^, which can be correlated to small quantities of SWCNTs. However, it is worth noting that SWCNTs benefit from signal enhancement due to resonance Raman scattering,^[66]^ and thus the intensity of their peaks in the shell clusters could over-represent their actual quantity with respect to PCL. Interestingly, the average Raman spectrum from the intermediate area features spectral characteristics from both SWCNTs and PCL polymer, indicating an area of heterogeneity at the core-shell interface. These results demonstrate that the highest density of the nanosensors reside in the core area of the fibers, while their Raman signal diminishes in the outward radial direction and the main component becomes the polymer shell. Although the spectral characteristics of the PEO polymer are not apparent in the Raman spectra of the three areas, they are presumably dominated by the enhanced Raman signal of the SWCNTs. Moreover, the Raman spectra of the PCL and PEO polymers show minimal overlap with the G-band and RBM peaks from SWCNTs (Figure S6a and b), indicating that the identified SWCNT signal contains no contribution from the other nanofiber components.

**Figure 3.**
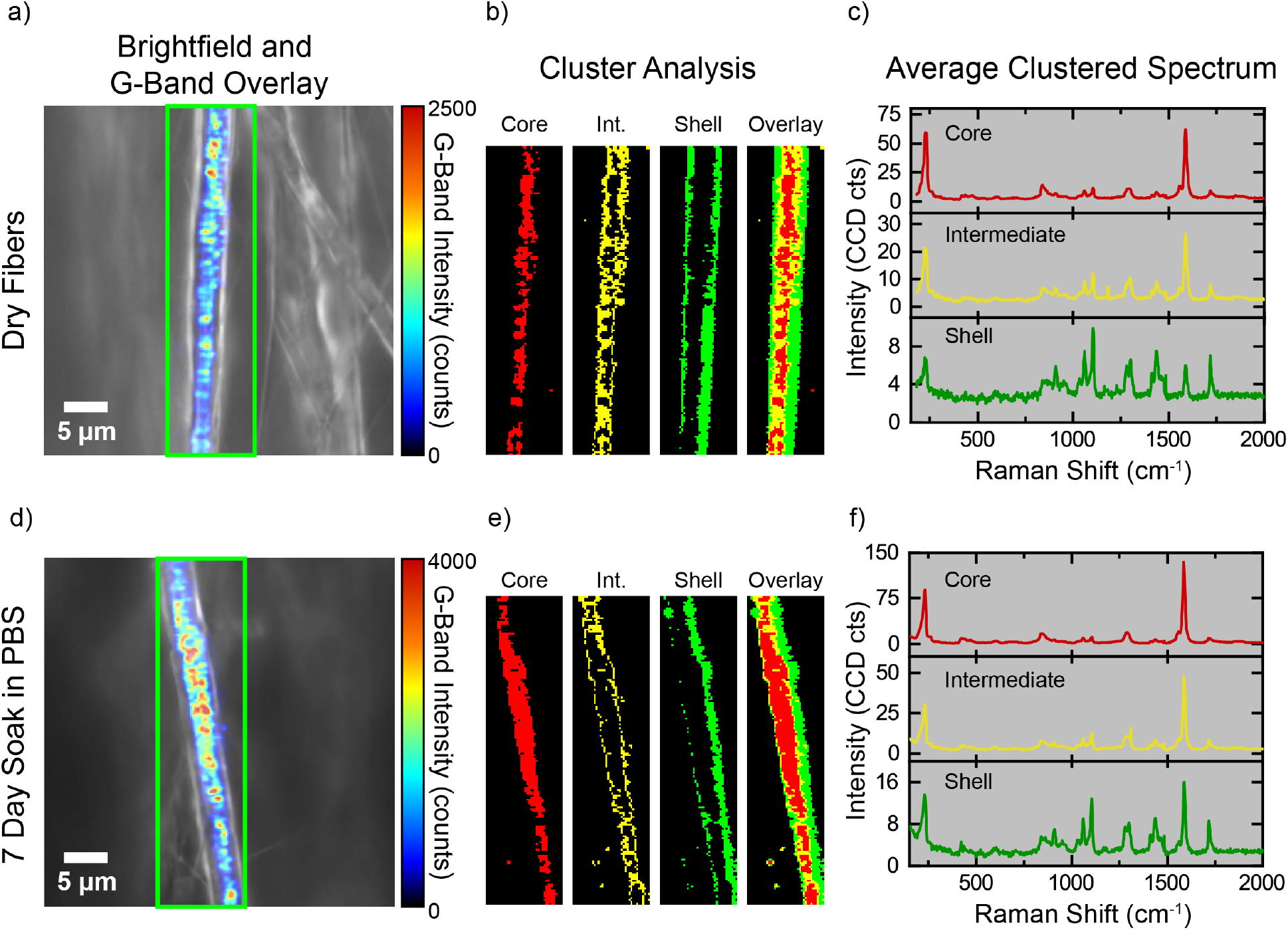
Confocal Raman microscopy of dry fibers and fibers soaked in PBS for 7 days. (a) and (d) The representative overlay of G-band intensity and brightfield images of dry fibers and fibers soaked in PBS for 7 days, respectively. (b) and (e) k-means clustering analysis of all spectra in each area scan, where *k* = 4 clusters (background cluster omitted from figure). (c) and (f) The average Raman spectra obtained from each cluster of (b) and (e).

To assess the ability of individual fibers to preserve the nanosensors over time, we soaked the fiber samples in phosphate buffered saline (PBS) solution and performed confocal Raman microscopy at different timepoints. Figure 3d and S7 indicate that the SWCNTs remain encapsulated within individual fibers after 7 days, without forming noticeable aggregates or any deformation in the fiber structure. Moreover, k-means clustering analysis identified the same spectral characteristics for the three identified areas (core, intermediate and shell) at different time points, further demonstrating that the fibers retain their entire structural integrity in an aqueous environment (Figure 3e, f, and S7b, c, e, f). We acknowledge that the ratios between the peaks in Raman spectra of particularly the shell vary among different time points, but this can be attributed to the slight heterogeneity among individual fibers as we have not imaged the same fiber at different time points.

To quantify the amount of the released nanosensors from a bulk fibrous matrix, the microfibrous textiles with thickness of ~0.7 mm were cut to 1 square inch pieces and soaked in PBS. We collected the PBS solutions over time for up to 21 days and acquired their Raman spectra (Figure S8). Comparing the Raman spectra of the released nanosensors with that of three standard samples with known concentrations (0.01, 0.1, 1 mg L^−1^), the lack of G-band signal indicates that the released nanosensors fall within the noise range of our Raman spectrometer. We conclude that over the course of at least 21 days, a negligible amount of the nanosensors are released from the fibers.

### 2.2. Ratiometric Peroxide Detection Using Optical Microfibrous Textiles

The HiPco SWCNTs contain multiple chiralities emitting NIR fluorescence in the range of 900-1400 nm (Figure S1b).^[39]^ As both chirality and DNA sequence determine the spectral responses of SWCNTs to their local environment,^41^ we hypothesized that certain chiralities of SWCNTs within the HiPco sample would differentially respond to hydrogen peroxide due to their differing bandgaps energies.^[67]^ We first acquired NIR hyperspectral fluorescence images of the fluorescent fibers containing (GT)_15_-SWCNTs (Movie S1) using the optical setup illustrated in Figure 4a. Figure 4c-e indicate the fluorescence intensity of three different chiralities in the same fiber sample, i.e. the (9,4), (8,6), and (8,7)-SWCNT chiralities.^[41]^ To test the environmental sensitivity of the optical fibers to peroxide, we exposed the bulk microfibrous samples (area: 0.5 mm^2^, thickness: ~0.7 mm) to various concentrations of peroxide and acquired fluorescence spectra after 24 hours, utilizing a probe NIR fluorescence spectrometer (Figure 4b), which enabled the resolution of the three mentioned chiralities. In agreement with previous studies,^[44, 47, 48]^ Figure 4f and S9 reveal that the three chiralities quench upon exposure to peroxide, however, the extent of quenching varies significantly among the chiralities. By normalizing each plot by its maximum intensity (i.e. the intensity of the (9,4) chirality), we observe that the normalized (8,6) and (8,7) peaks monotonically intensify with increasing peroxide concentration, illustrating that a ratiometric signal can be obtained for peroxide detection (Figure 4g). We selected the (8,7) / (9,4) intensity ratio to further calibrate a biosensor for peroxide detection as it appears to be more sensitive to peroxide concentrations compared to the (8,6) / (9,4) ratio.

**Figure 4.**
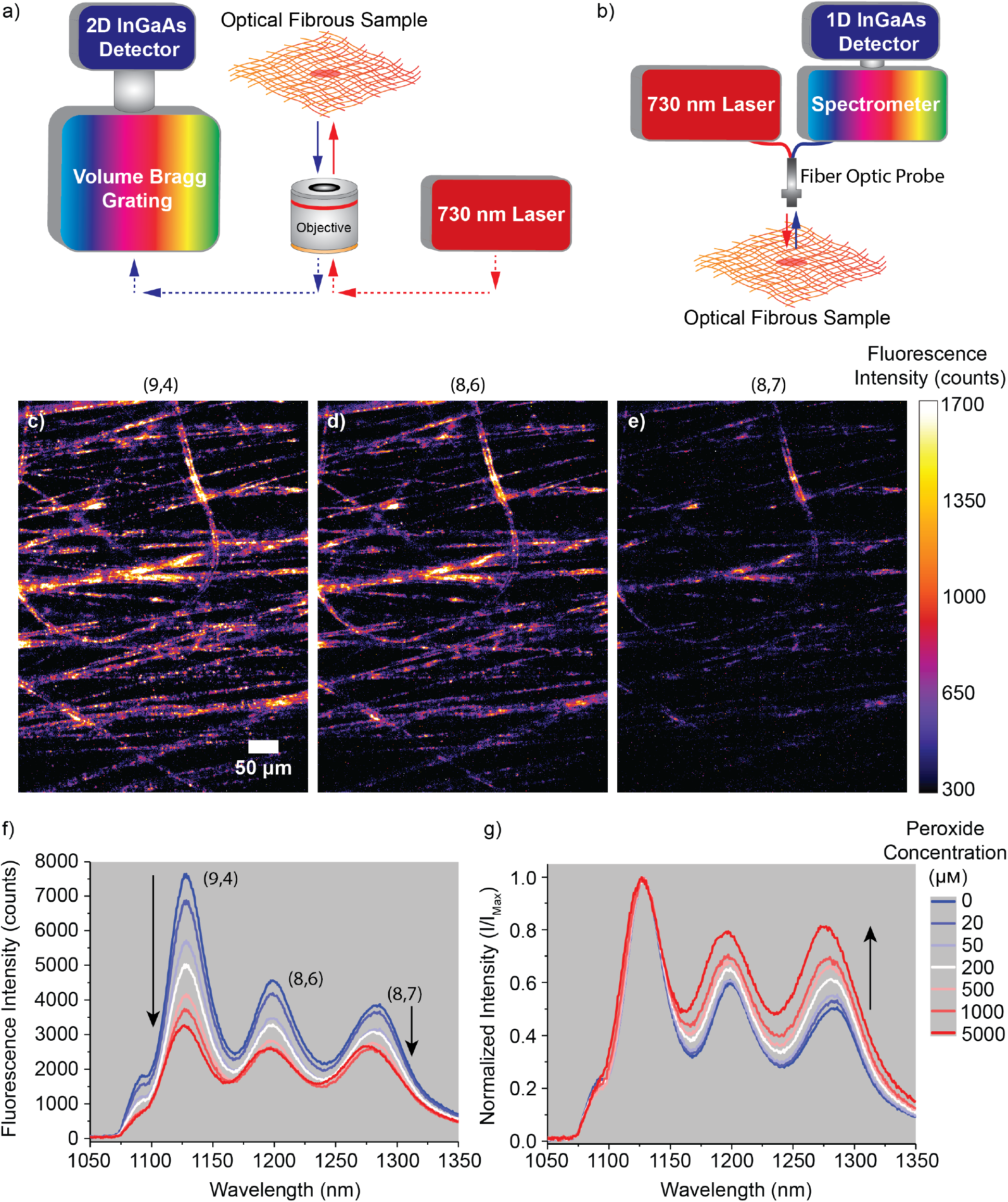
(a) The schematic of the optical setup utilized to obtain the hyperspectral NIR fluorescence intensity images from the microfibers. (b) The schematic of the probe NIR fluorescence spectrometer utilized to obtain the fluorescence spectra from bulk microfibrous samples. (c) (9,4), (d) (8,6) and (e) (8,7) chirality intensity images, obtained using the system shown in part a. (e) The fluorescent spectra of the microfibrous samples exposed to various peroxide concentrations, obtained from the system shown in part b. (f) The fluorescent spectra shown in part d were normalized by their max intensity.

### 2.3. Calibrating the Microfibrous Textiles for Peroxide Detection

To obtain a calibration curve for aqueous peroxide detection, we exposed the initially dry microfibrous samples to various concentrations of peroxide ranging from 1 μM to 5 mM and monitored the samples over time for up to 72 hours. Figure 5a demonstrates that all samples produced the same initial ratiometric signal, clarifying the reproducibility of our method for fabricating bulk samples of optical fibers encapsulating nanosensors. After 24 hours, a concentration-dependent ratiometric signal was obtained from the samples. While 0 to 5 μM peroxide gave no significant change in the ratiometric signal, it monotonically increased with peroxide concentration in the range of 5 μM to 5 mM and could be fit to a linear function with R^2^ = 0.99 on a log-log scale of signal vs. concentration (Figure 5a and Table S1). When examined at 48 and 72 hours after the addition of peroxide, the ratiometric signal systematically increases while maintaining the trend in the calibration curve. The data can still be fit to a linear function in the range of 5 μM to 5 mM. To explain the time dependency of the calibration curves, we propose that the noncovalently wrapped DNA adopts more compact conformations on the SWCNT surfaces over time as they progressively interact with ions, mainly sodium, in the PBS.^[68]^ This rearrangement alters the fluorescence intensity in a chirality-dependent fashion, and thus the ratio of the peaks over time, regardless of peroxide concentration.^[68]^ Moreover, the diffusion of the peroxide molecules through the pores in the 3D matrix and through the free spaces in between the polymer chains on the shell is a time-dependent process, so it will result in further fluorescence quenching over time. These two phenomena governing the temporal dependence of the peroxide calibration curve are depicted in Figure 5b.

**Figure 5.**
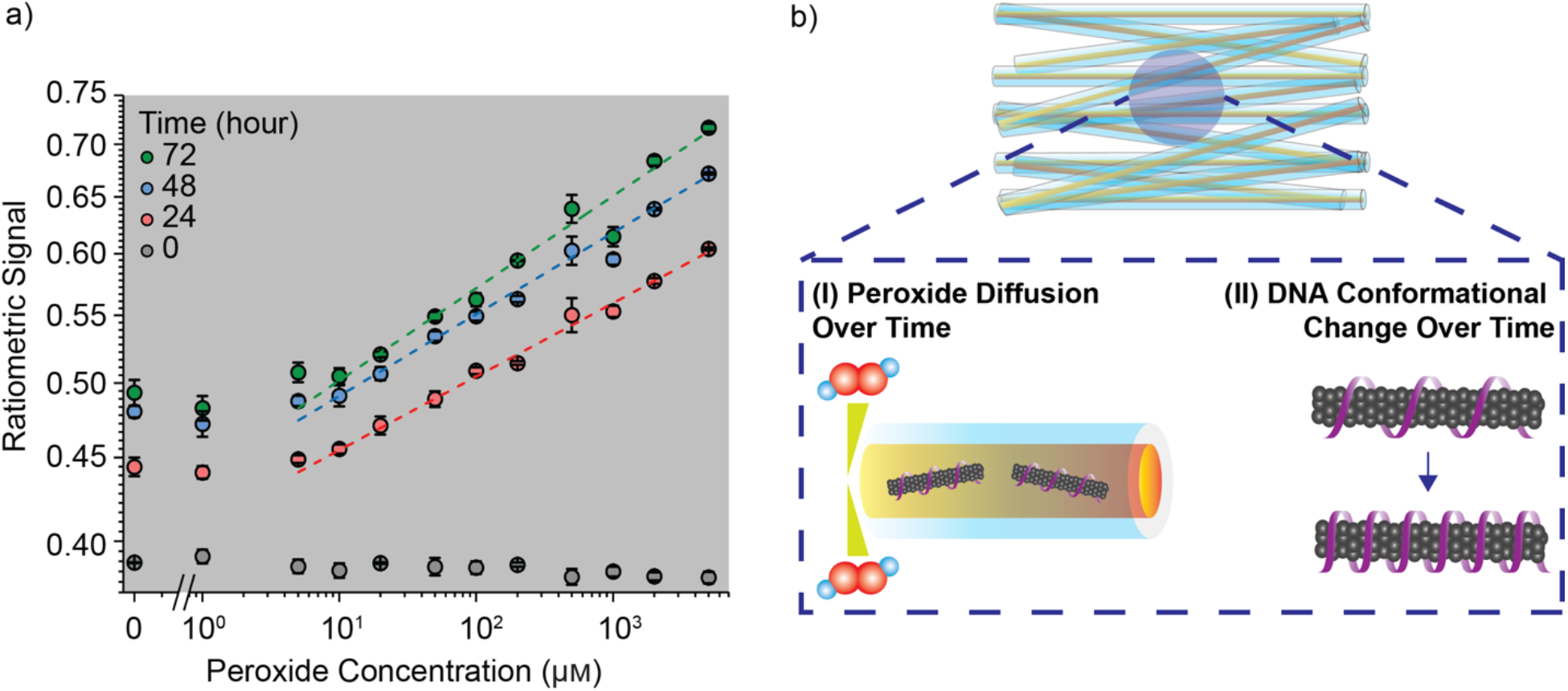
(a) Calibration curve showing the ratiometric signal, i.e. (8,7) intensity divided by (9,4) intensity, as a function of peroxide concentration at three different time points. Mean values were obtained by adding each peroxide concentration to three different samples (*n* = 3), and error bars represent the standard deviation. (b) Two phenomena governing the temporal dependence of the peroxide calibration curve; (I) The diffusion of peroxide molecules through the 3D matrix of fibers and through the shell to reach to the ssDNA-SWCNTs in the core of the individual fibers. (II) Due to the ionic strength of the surrounding environment, the ssDNA on the surface of nanotubes undergoes a conformational change over time and forms a more compact wrapping around the nanotubes. As a result of these two phenomena, the NIR fluorescence of SWCNTs and the ratiometric signal are altered over time.

Therefore, in order to design a wearable fibrous device for continuous peroxide monitoring, we propose a two-dimensional calibration curve where the ratiometric signal is a function of both analyte concentration and time. The heatmap in Figure 6a demonstrates the ratiometric signal as a function of the tested concentrations of peroxide and time points. We plotted the ratiometric signal as a function of time for all concentrations (Figure 6b). Interestingly, the data for all examined peroxide concentrations could be fit to single exponential association functions (Equation 1 and Figure S10 and S11) where the offset (*R_0_*) and time constant (*τ*) of the single exponential showed a narrow distribution with small standard errors of the means (Figure S12 and Table S2). The lack of dependence on peroxide concentration found within the fitted time constants, including the dataset of zero added peroxide, suggests the dominant physical process being modeled is that of the DNA rearrangement on the SWCNT surface. In contrast, the pre-exponential factors (*A*) displayed a concentration-dependent trend which could be fit to a power function (Equation 2, Figure 6c and S13, Table S3). By incorporating Equation 2 into Equation 1, we obtained a two-input calibration function where the ratiometric signal could be expressed as a function of both peroxide concentration and time (Equation 3, Figure 6d). Finally, to examine the reversibility of the platform, we exposed a sample to peroxide (200 μM) and then washed it with PBS. As shown in Figure S14, we observed that the ratiometric signal increased 10 minutes after exposure to the peroxide and then completely reverts back to the original signal 50 minutes after removing it.

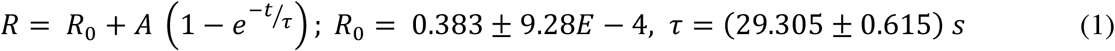

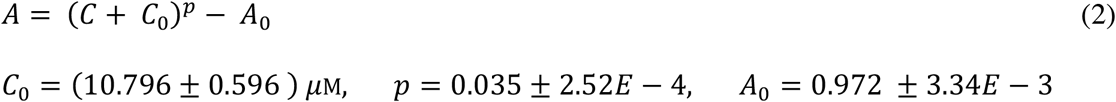

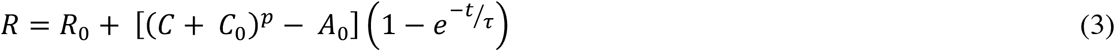

**Figure 6.**
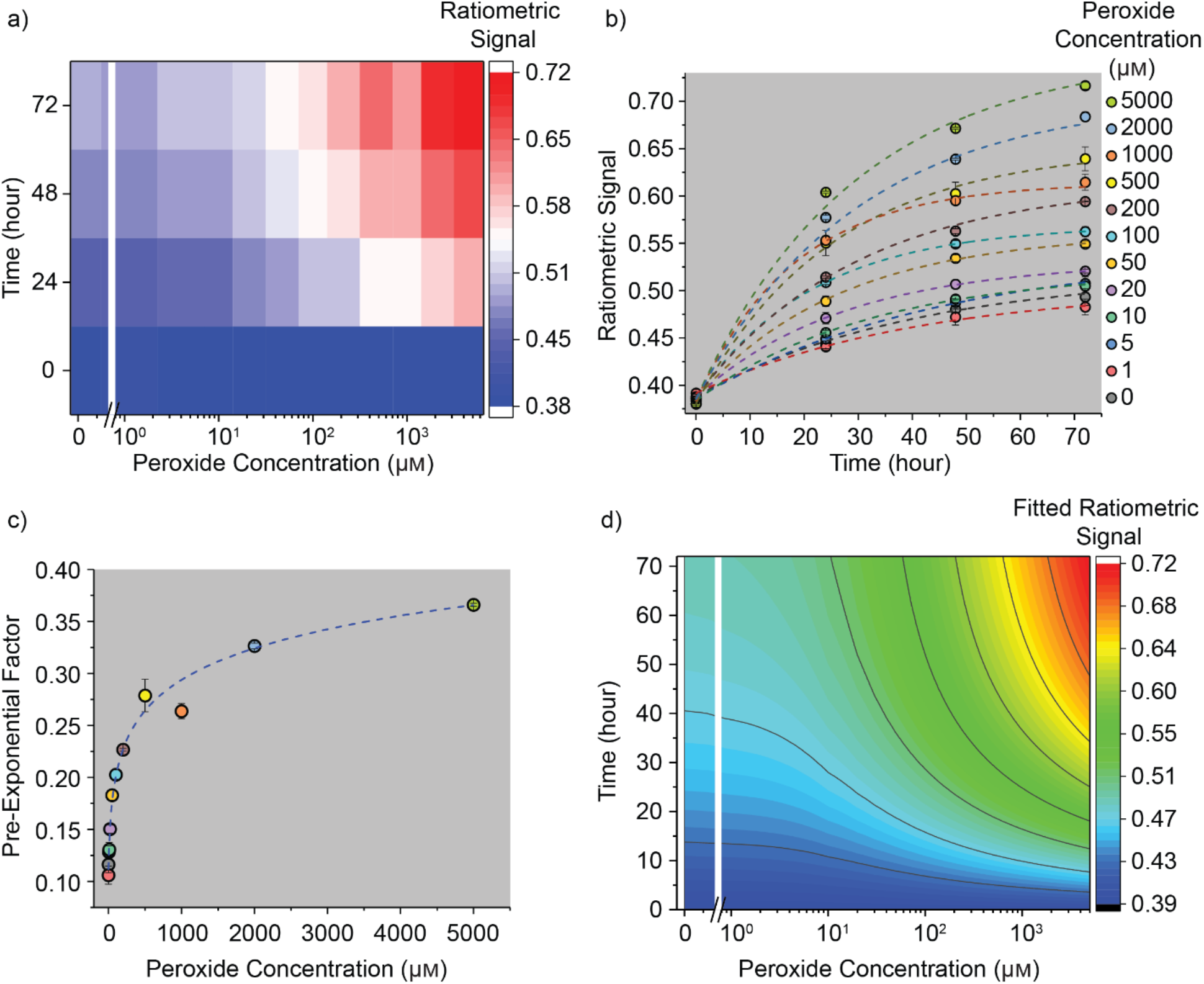
Calibrating the optical microfibrous textiles for continuous peroxide detection. (a) Two-dimensional heat-map illustrating the ratiometric signal as a function of both concentration and time. (b) Ratiometric signal as a function of time for each peroxide concentration. The dashed lines indicate single exponential association fits. (c) Pre-exponential factors extracted from the single exponential association fits and plotted as a function of peroxide concentration. The dashed line indicates a power function fit. (d) Contour plot demonstrating a two-dimensional calibration curve where the fitted ratiometric signal is a function of both time and concentration. Mean values were obtained by adding each peroxide concentration to three different samples (*n* = 3), and error bars represent the standard deviation.

In contrast to an average concentration reported by a single readout, the ability to spatially resolve peroxide concentrations will enable an end-user the ability to map peroxide on the surface of a wound utilizing an NIR camera with the appropriate filters. We exposed the microfibrous samples to different concentrations of peroxide and acquired hyperspectral fluorescence images from the surface of the samples under 5X magnification using the setup shown in Figure 4a. By dividing the maximum intensity images of the (8,7)-SWCNT by the (9,4)-SWCNT, we created maps where each pixel represented a ratiometric signal (Figure 7). At time zero, the maps for all concentrations of peroxide were dominated by blue pixels (low ratiometric signal). After 24-hour exposure to peroxide, the pixel colors among the maps diverge and are dominated by yellow color at low concentrations and red color at high concentrations. This demonstrates the potential of employing our optical fibrous platform to quantitatively image the surface of a wound in a label-free manner.

**Figure 7.**
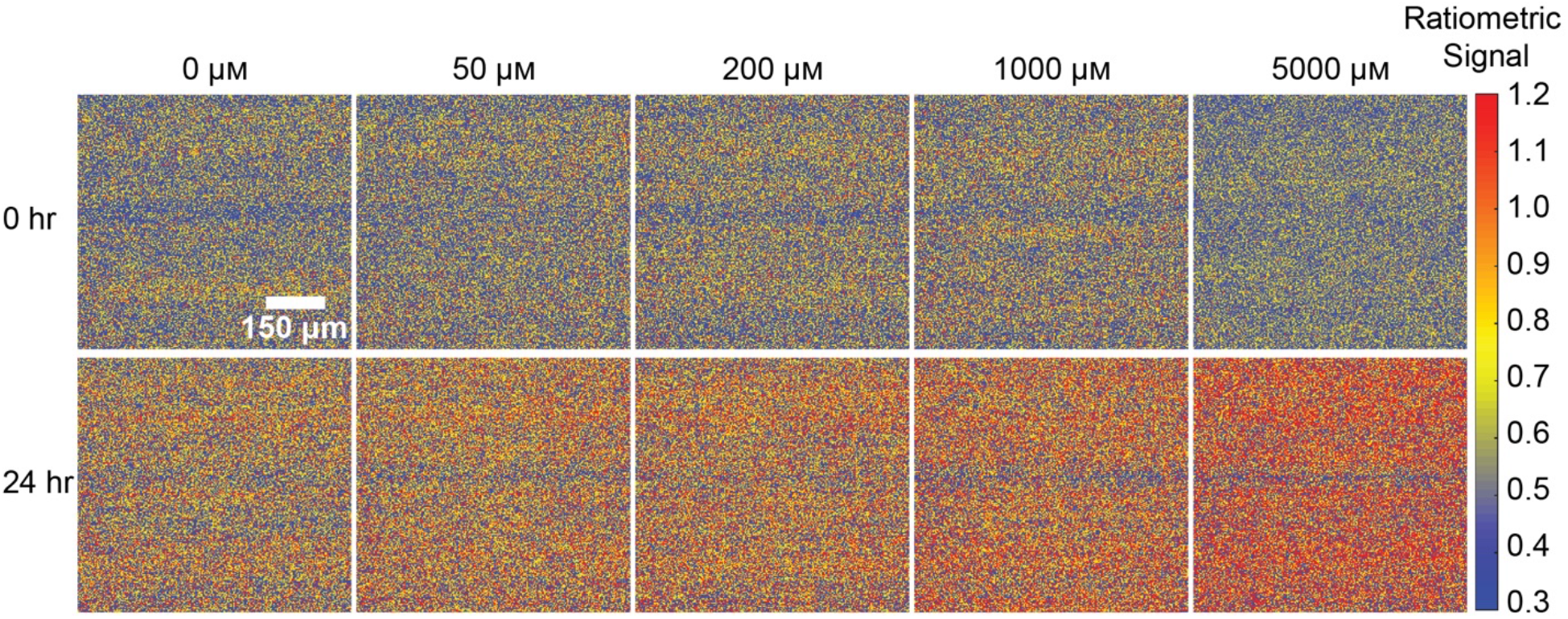
Spatially detecting peroxide. The maps were created by acquiring NIR fluorescence hyperspectral images and dividing the max intensity image of the (8,7)-SWCNT by the max intensity image of the (9,4) chirality SWCNT. Since the data were acquired using a 5X objective, the individual fibers cannot be observed.

To illustrate the potential of our optical fibrous platform as a smart wound dressing for *in situ* monitoring of peroxide, we attached the fibrous samples onto a commercial wound bandage (Figure 8a). By attaching the samples to the complete bandage (adhesive material + adsorbent pad) or only to the adhesive material of the bandage, the fluorescence of SWCNTs was still detectable by our probe spectrometer through the bandage and with a high signal to noise ratio for each peak (Figure 8b and S15). The drop in the signal compared to the control is presumably due to the polymers in the bandage that absorb a portion of the excitation light and/or emitted fluorescence from the SWCNTs. Additionally, the combination of sample plus adhesive did not significantly alter the ratiometric signal even after 7 days of soaking in PBS (Figure 8c). Finally, Figure 8d and Movie S2 demonstrate the feasibility of a real-time wireless wound screening utilizing our flexible optical fibrous platform attached onto the commercial bandage.

**Figure 8.**
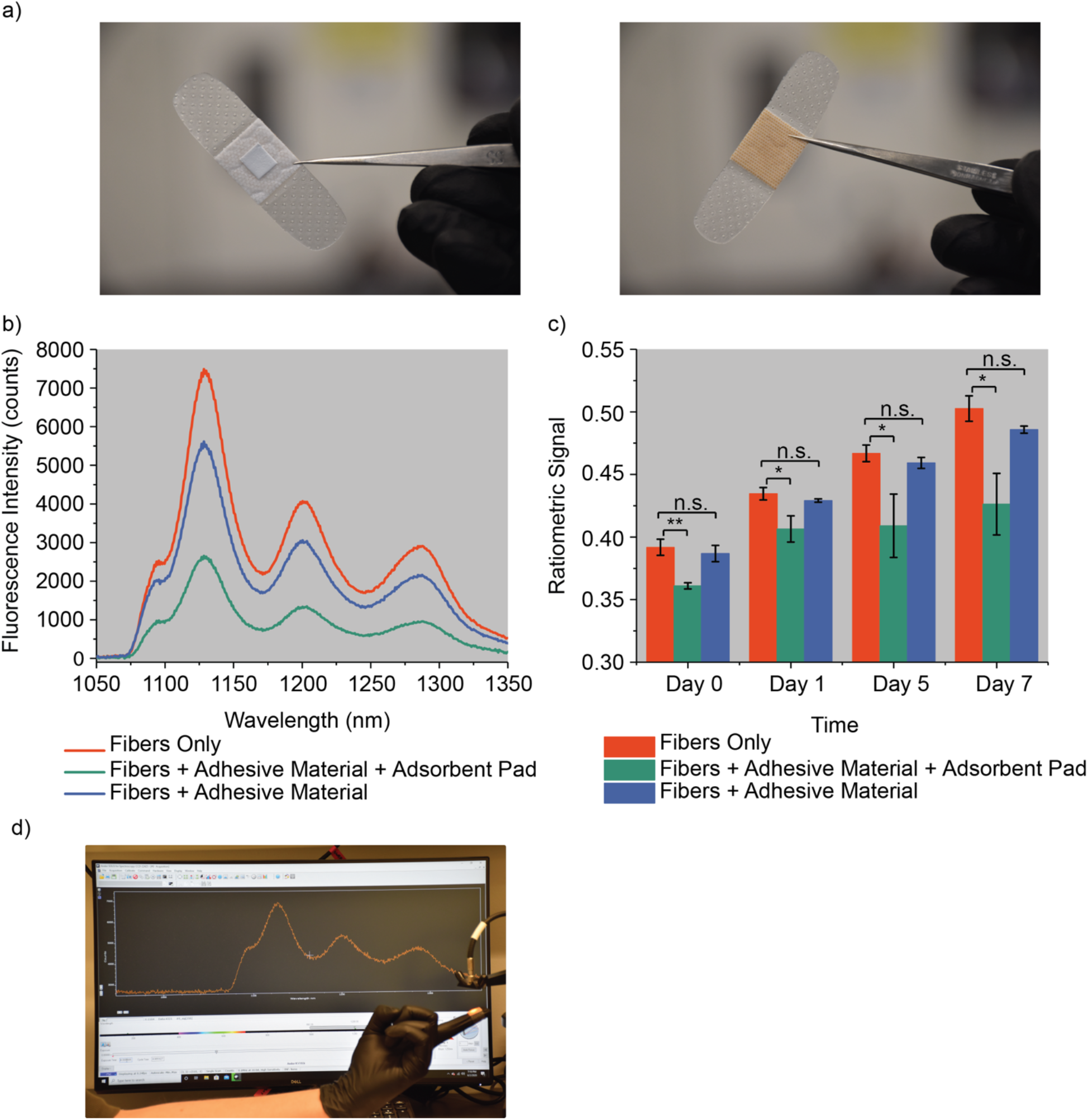
(a) Integrating the optical fibrous samples into a commercial wound bandage. (b) Comparison of the fluorescence spectra of microfibers alone, through adhesive bandage material, or through both adhesive material plus an adsorbent pad (complete bandage). (c) Comparison of the ratiometric signals of the three conditions mentioned in part b, after being soaked in PBS over time. Mean values were obtained by acquiring the fluorescence spectra from three different samples (*n* = 3) per each condition. The error bars represent standard deviation. Two-sample t-tests were performed on the data (*, *p* < 0.05, **, *p* < 0.01). (d) Real-time wireless fluorescence spectra readout utilizing the flexible optical microfibers attached onto a commercial bandage.

## 3. Conclusions

Multi-compartment smart wound dressings have attracted a substantial interest in the past few years due to their potential for enabling simultaneous wound monitoring and healing.^[51, 52]^ A smart wound dressing usually integrates multiple layers into a single flexible biomaterial platform.^[69]^ A therapeutic layer can be designed to enable wound healing by incorporating antibiotics, growth factors, etc., into a 3D biocompatible scaffold such as hydrogels and microfibers. Moreover, a sensing layer would enable continuous monitoring of multiple biomarkers in the wound environment. In this work, we employed a one-step coaxial electrospinning process to encapsulate ssDNA-SWCNT nanosensors into individual microfibers to fabricate wearable optical microfibrous textiles, as a sensing layer, for monitoring oxidative stress in wounds. Utilizing confocal Raman microscopy of individual fibers over time, we uncovered that the SWCNT nanosensors are preserved inside of the fibers over time and that a negligible quantity of the nanosensors are released from a 3D fibrous matrix after 21 days. As multiple nanotube chiralities in the HiPco sample respond differentially to peroxide molecules, we designed an optical wearable platform to ratiometrically detect peroxide at physiologically relevant concentrations. Utilizing this flexible optical microfibrous material, the wireless detection of peroxide was demonstrated. In addition to a single real-time readout, we demonstrated the potential of our platform for spatially resolving the peroxide concentrations on a wound surface using an InGaAs camera, without the need to use any additional marker. We finally integrated our microfibrous textiles onto a commercial wound bandage and indicated the compatibility of this platform with existing wound dressings.

## 4. Materials and Methods

### ssDNA-SWCNT Nanosensor Preparation

Raw single-walled carbon nanotubes produced by the HiPco process (1 mg, Nanointegris) were added to desalted (GT)_15_ oligonucleotide (2 mg, Integrated DNA Technologies) in a microcentrifuge tube with NaCl solution (1 mL, 0.1 M, Sigma-Aldrich). The mixture was then ultrasonicated using a 1/8″ tapered microtip (Sonics Vibracell; Sonics & Materials) for 30 min at 40% amplitude, with an average power output of 8 W, in a 0 °C temperature-controlled microcentrifuge tube holder. After sonication, the dispersion was ultracentrifuged twice (Sorvall Discovery M120 SE) for 30 min at 250 000*×g*, and the top 80% of the supernatant was extracted. The resultant dispersion was filtered using 100 kDa Amicon centrifuge filters (Millipore) to remove free ssDNA. A UV/vis/NIR spectrophotometer (Jasco, Tokyo, Japan) was utilized to determine the concentration using the extinction coefficient of A_910_ = 0.02554 L mg^−1^ cm^−1^.^[32, 33]^

### Preparation of Core and Shell Polymer Solutions

A 4 wt.% poly(ethylene oxide) (PEO, M_v_ = 900,000 g mol^−1^, Sigma-Aldrich) solution was prepared by dissolving PEO in DI water and stirring the solution overnight on a hotplate set to 48 °C. A concentrated ssDNA-SWCNT dispersion (~400-500 mg L^−1^) was prepared by filtering out the as-prepared ssDNA-SWCNT dispersion using an Amicon filter (100 kDa) and resuspending it in a lower volume of NaCl solution (0.1 M). The concentrated dispersion was then diluted in the resultant PEO solution to obtain a homogenous nanotube concentration of 10 mg L^−1^. Because of the high ssDNA-SWCNT concentration, the final concentration of the PEO solution was not significantly altered by adding ssDNA-SWCNT dispersion to it. Polycaprolactone (PCL, M_w_ = 70,000 g mol^−1^, Scientific Polymer Products, Inc.) was dissolved in a mixture of chloroform and dimethylformamide (DMF) with the volume ratio of 80:20, by stirring the solution for 6 hours at room temperature, to obtain a final PCL concentration of 13 wt.%.

### Fabrication of Electrospun Optical Microfibrous Textiles

A one-step co-axial electrospinning process was used to produce core-shell fibers. Figure 1b illustrates the schematic of the experimental setup. A customized core-shell needle (Rame-hart Instrument co.) with two separate inlets was built by placing a 24 Gauge needle inside of a 15 Gauge needle. The inlets of the needle were connected to two syringes filled with the polymer solutions and placed on a syringe pump capable of controlling the flow rates separately. The flow rates of the core and shell solutions were set to 0.3 and 2 mL h^−1^, respectively. A high voltage supply was connected to the tip of the needle and the rotating collector was grounded. The working distance between the needle and collector was set to 12 cm. To fabricate bulk fibrous textiles with a thickness of ~0.7 mm, the fibers were continuously collected on the metal collector for 7 hours. To prepare samples for NIR and confocal Raman microscopy, microscope coverslips were taped to the surface of the collector and a thin layer the fibers were collected on the coverslips for 10 minutes.

### Near-Infrared Fluorescence Microscopy of the Fibers

As described previously,^[41, 70]^ a near-infrared hyperspectral fluorescence microscope was used to acquire fluorescence images and hyperspectral cubes from a thin layer of fibers collected on a microscope coverslip. In short, a continuous 730 nm diode laser with 2.5 W output power was injected into a multimode fiber to produce an excitation source, which was reflected on the sample stage of an Olympus IX-73 inverted microscope equipped with a 20X LCPlan N, 20x/0.45 IR objective (Olympus, U.S.A.). To generate spectral image stacks (cubes), the emission was passed through a volume Bragg grating and collected with a 2D InGaAs array detector (Photon Etc.) with a spectral resolution of 4 nm. A background subtraction was performed using a custom MATLAB code. The background-subtracted images and hyperspectral cubes were processed and extracted using the Fiji software.

### Scanning Electron Microscopy of the Fibers

The samples were sputtered with gold prior to imaging. The SEM images were taken using a Zeiss Sigma VP field emission scanning electron microscope (FE-SEM), with an InLens detector and an accelerating voltage of 3.00 kV.

### Confocal-Raman Microscopy

A thin layer of fibers collected on a microscope coverslip was imaged with a WiTec Alpha300 R confocal-Raman microscope (WiTec, Germany) equipped with a Zeiss EC Epiplan-Neofluar 100×/0.9 air objective, a 785 nm laser source set to 35 mW sample power, and collected with a UHTS 300 spectrograph (300 lines/mm grating) coupled with an Andor DR32400 CCD detector (−61 °C, 1650 × 200 pixels). 10 × 40 μm areas were scanned, and spectra were obtained in 0.25 × 0.25 μm intervals with 0.4 s integration time to construct hyperspectral images of individual fibers. Global background subtraction, cosmic-ray removal, and k-means cluster analysis were performed on each scan using WiTec Project 5.2 software. G-band intensity images were constructed by integrating the spectrum of each SWCNT-containing pixel from 1575 to 1605 cm^−1^ using custom Matlab codes.

### Quantifying the Amount of the Released Nanosensor Using Solution-Based Raman Spectroscopy

1 square inch pieces of the bulk fibrous samples with thickness of ~ 0.7 mm were soaked in 3 mL of PBS (1X) over time. The PBS was collected at different time points (1 hour, 24 hours, 48 hours, 72 hours, 7 Days, 14 Days and 21 Days) and replaced with 3 mL of fresh PBS. The collected samples were placed into glass vials and Raman spectra were obtained using an inverted WiTec Alpha300 R confocal Raman microscope (WiTec, Germany) equipped with a Zeiss Epiplan-Neofluar 10×/0.25 objective, a 785 nm laser source set to 50 mW sample power, and collected with a UHTS 300 spectrograph (300 lines/mm grating) coupled with an Andor DR32400 CCD detector (1650 × 200 pixels). Each spectrum was averaged 5 times with 5 s integration time. Background subtraction was performed on all data using Witec Project 5.2 software.

### Real-Time Near-Infrared Fluorescence Spectroscopy of Bulk Microfibrous Samples

Bulk fibrous samples with thickness of ~0.7 mm were placed into plastic Petri dishes and 3 mL of the peroxide solution diluted to different concentrations in PBS (1X) was added to the samples. The NIR fluorescence spectra was acquired from each sample at 24, 48 and 72 hours. Individual NIR fluorescence spectra from the bulk samples were obtained using a custom-built preclinical fiberoptic probe spectroscopy system described in previous studies.^[34, 35]^ In summary, a continuous-wave 1320-mW 730-nm laser (CNI lasers) was injected into a bifurcated fiber optic reflection probe bundle. The bundle consisted of a 200-mm, 0.22 numerical aperture (NA) fiber optic cable for sample excitation located in the center of six 200-mm, 0.22 NA fibers for collection. Long-pass filters were used to block emission below 1100 nm. The light was focused into a 193-mm focal length Czerny-Turner spectrograph (Kymera 193i, Andor) with the slit width set at 410 mm. Light was dispersed by an 1501/mm grating with blaze wavelength of 1250 nm and collected with an iDus InGaAs camera (Andor). The distance between the laser probe tip and the sample was set to 2.2 cm. A custom MATLAB code was used to perform background subtraction on the acquired fluorescence spectra.

### Statistical Analysis

All curve fittings, statistical measurements and analyses were performed in OriginPro 2016. The two-sample t-tests were performed under the null hypothesis.

## Table of Contents Figure

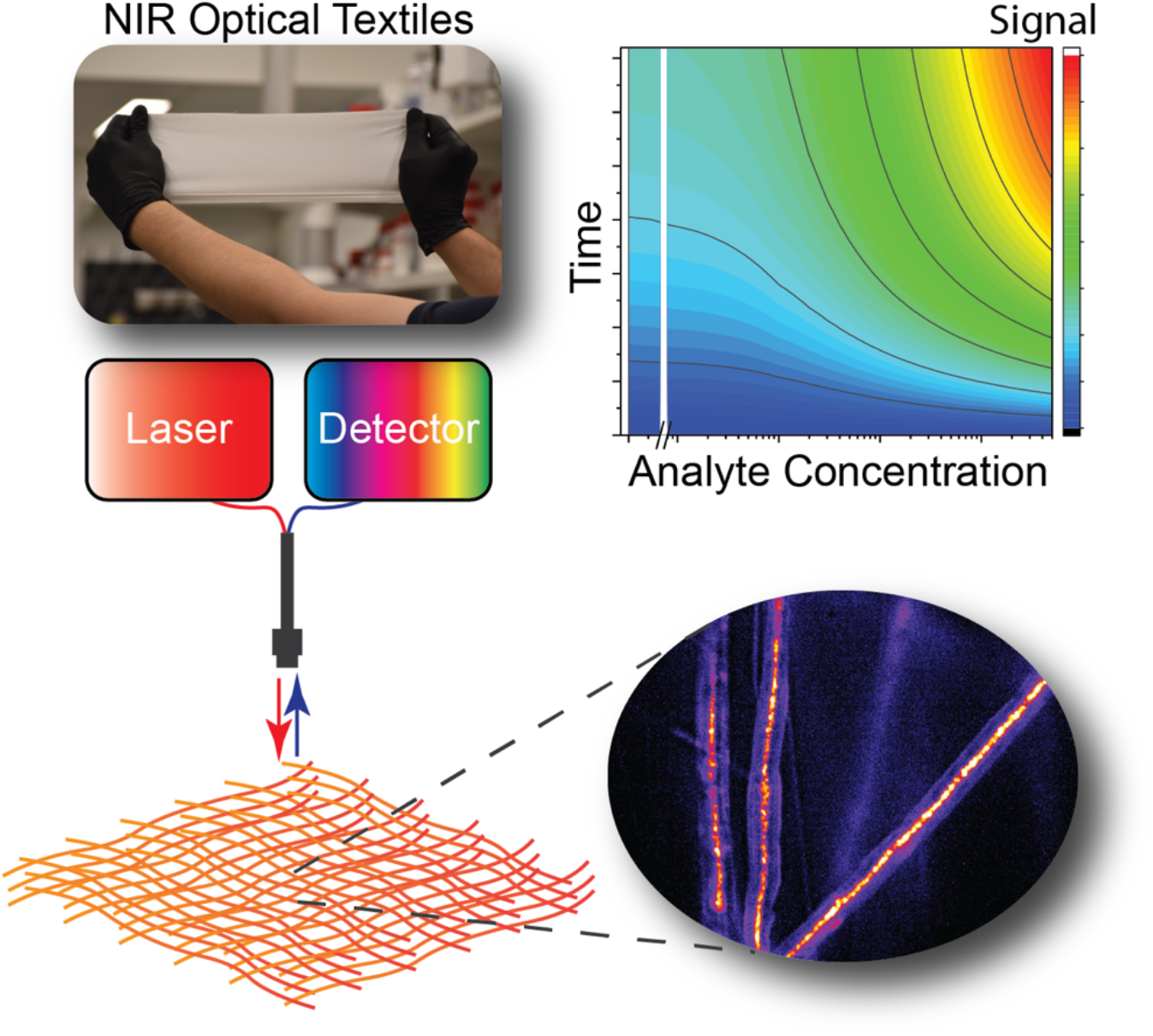

## Supporting Information

Supporting Information is available online.

## Acknowledgments

This work was supported by National Science Foundation CAREER Award #1844536, the RI-INBRE Early Career Development Award Grant P20GM103430 from the National Institute of General Medical Sciences of the National Institutes of Health, the Rhode Island Foundation−Medical Research Fund, and the URI College of Engineering. The confocal Raman data and SEM images were acquired at the RI Consortium for Nanoscience and Nanotechnology, a URI College of Engineering core facility partially funded by the National Science Foundation EPSCoR, Cooperative Agreement #OIA-1655221. We acknowledge I. Andreu for the help in taking SEM images. We acknowledge P. V. Jena for designing, building and installing the custom preclinical fiberoptic probe spectroscopy system.

## References

[1] C. Nathan, A. Cunningham-Bussel, Nat. Rev. Immunol. 2013, 13, 349.

[2] B. C. Dickinson, C. J. Chang, Nat. Chem. Biol. 2011, 7, 504.

[3] C. C. Winterbourn, Nat. Chem. Biol. 2008, 4, 278.

[4] F. C. Fang, Nat. Rev. Microbiol. 2004, 2, 820.

[5] D. K. Singh, P. Winocour, K. Farrington, Nat. Rev. Endocrinol. 2011, 7, 176.

[6] C. Gorrini, I. S. Harris, T. W. Mak, Nat. Rev. Drug Discovery 2013, 12, 931.

[7] S. Bisht, M. Faiq, M. Tolahunase, R. Dada, Nat. Rev. Urol. 2017, 14, 470.

[8] P. Niethammer, C. Grabher, A. T. Look, T. J. Mitchison, Nature 2009, 459, 996.

[9] Y. Huo, W.-Y. Qiu, Q. Pan, Y.-F. Yao, K. Xing, M. F. Lou, Exp. Eye Res. 2009, 89, 876.

[10] N. Suzuki, R. Mittler, Free Radical Biol. Med. 2012, 53, 2269.

[11] C. Dunnill, T. Patton, J. Brennan, J. Barrett, M. Dryden, J. Cooke, D. Leaper, N. T. Georgopoulos, Int. Wound J. 2017, 14, 89.

[12] M. P. Brynildsen, J. A. Winkler, C. S. Spina, I. C. MacDonald, J. J. Collins, Nat. Biotechnol. 2013, 31, 160.

[13] R. Zhao, H. Liang, E. Clarke, C. Jackson, M. Xue, Int. J. Mol. Sci. 2016, 17, 2085.

[14] R. Moseley, J. R. Hilton, R. J. Waddington, K. G. Harding, P. Stephens, D. W. Thomas, Wound Repair Regen. 2004, 12, 419.

[15] G. Zhao, M. L. Usui, S. I. Lippman, G. A. James, P. S. Stewart, P. Fleckman, J. E. Olerud, Adv. Wound Care 2013, 2, 389.

[16] S. Dhall, D. C. Do, M. Garcia, J. Kim, S. H. Mirebrahim, J. Lyubovitsky, S. Lonardi, E. A. Nothnagel, N. Schiller, M. Martins-Green, J. Diabetes Res. 2014, 2014.

[17] J. H. Kim, B. Yang, A. Tedesco, E. G. D. Lebig, P. M. Ruegger, K. Xu, J. Borneman, M. Martins-Green, Sci. Rep. 2019, 9, 1.

[18] A. E. K. Loo, Y. T. Wong, R. Ho, M. Wasser, T. Du, W. T. Ng, B. Halliwell, PLoS One 2012, 7, e49215.

[19] S. Roy, S. Khanna, K. Nallu, T. K. Hunt, C. K. Sen, Mol. Ther. 2006, 13, 211.

[20] M. Schäfer, S. Werner, Pharmacol. Res. 2008, 58, 165.

[21] M. Cano Sanchez, S. Lancel, E. Boulanger, R. Neviere, Antioxidants 2018, 7, 98.

[22] P. M. Abuja, R. Albertini, Clin. Chim. Acta 2001, 306, 1.

[23] T. J. James, M. A. Hughes, G. W. Cherry, R. P. Taylor, Wound Repair Regen. 2003, 11, 172.

[24] M. Mohammadniaei, J. Yoon, T. Lee, B. G. Bharate, J. Jo, D. Lee, J. W. Choi, Small 2018, 14, 1703970.

[25] B. Ma, C. Kong, X. Hu, K. Liu, Q. Huang, J. Lv, W. Lu, X. Zhang, Z. Yang, S. Yang, Biosens. Bioelectron. 2018, 106, 29.

[26] H. Jin, D. A. Heller, M. Kalbacova, J.-H. Kim, J. Zhang, A. A. Boghossian, N. Maheshri, M. S. Strano, Nat. Nanotechnol. 2010, 5, 302.

[27] D. A. Heller, H. Jin, B. M. Martinez, D. Patel, B. M. Miller, T.-K. Yeung, P. V. Jena, C. Höbartner, T. Ha, S. K. Silverman, Nat. Nanotechnol. 2009, 4, 114.

[28] W.-K. Oh, Y. S. Jeong, S. Kim, J. Jang, Acs Nano 2012, 6, 8516.

[29] Y. G. Ermakova, D. S. Bilan, M. E. Matlashov, N. M. Mishina, K. N. Markvicheva, O. M. Subach, F. V. Subach, I. Bogeski, M. Hoth, G. Enikolopov, Nat. Commun. 2014, 5, 1.

[30] V. V. Belousov, A. F. Fradkov, K. A. Lukyanov, D. B. Staroverov, K. S. Shakhbazov, A. V. Terskikh, S. Lukyanov, Nat. Methods 2006, 3, 281.

[31] Y. Fujikawa, L. P. Roma, M. C. Sobotta, A. J. Rose, M. B. Diaz, G. Locatelli, M. O. Breckwoldt, T. Misgeld, M. Kerschensteiner, S. Herzig, Sci. Signaling 2016, 9, rs1.

[32] M. M. Safaee, M. Gravely, A. Lamothe, M. McSweeney, D. Roxbury, Sci. Rep. 2019, 9, 1.

[33] M. M. Safaee, M. Gravely, C. Rocchio, M. Simmeth, D. Roxbury, ACS Appl. Mater. Interfaces 2018, 11, 2225.

[34] J. Budhathoki-Uprety, J. Shah, J. A. Korsen, A. E. Wayne, T. V. Galassi, J. R. Cohen, J. D. Harvey, P. V. Jena, L. V. Ramanathan, E. A. Jaimes, Nat. Commun. 2019, 10, 1.

[35] T. V. Galassi, P. V. Jena, J. Shah, G. Ao, E. Molitor, Y. Bram, A. Frankel, J. Park, J. Jessurun, D. S. Ory, Sci. Transl. Med. 2018, 10, eaar2680.

[36] J. D. Harvey, R. M. Williams, K. M. Tully, H. A. Baker, Y. Shamay, D. A. Heller, Nano Lett. 2019, 19, 4343.

[37] C.-W. Lin, S. M. Bachilo, Y. Zheng, U. Tsedev, S. Huang, R. B. Weisman, A. M. Belcher, Nat. Commun. 2019, 10, 1.

[38] A. G. Godin, A. Setaro, M. Gandil, R. Haag, M. Adeli, S. Reich, L. Cognet, Sci. Adv. 2019, 5, eaax1166.

[39] S. M. Bachilo, M. S. Strano, C. Kittrell, R. H. Hauge, R. E. Smalley, R. B. Weisman, Science 2002, 298, 2361.

[40] P. V. Jena, D. Roxbury, T. V. Galassi, L. Akkari, C. P. Horoszko, D. B. Iaea, J. Budhathoki-Uprety, N. Pipalia, A. S. Haka, J. D. Harvey, ACS nano 2017, 11, 10689.

[41] M. Gravely, M. M. Safaee, D. Roxbury, Nano Lett. 2019, 19, 6203.

[42] P. V. Jena, M. M. Safaee, D. A. Heller, D. Roxbury, ACS Appl. Mater. Interfaces 2017, 9, 21397.

[43] R. M. Williams, C. Lee, T. V. Galassi, J. D. Harvey, R. Leicher, M. Sirenko, M. A. Dorso, J. Shah, N. Olvera, F. Dao, Sci. Adv. 2018, 4, eaaq1090.

[44] T. T. S. Lew, V. B. Koman, K. S. Silmore, J. S. Seo, P. Gordiichuk, S.-Y. Kwak, M. Park, M. C.-Y. Ang, D. T. Khong, M. A. Lee, Nat. Plants 2020, 6, 404.

[45] M. Dinarvand, E. Neubert, D. Meyer, G. Selvaggio, F. A. Mann, L. Erpenbeck, S. Kruss, Nano Lett. 2019, 19, 6604.

[46] A. G. Godin, J. A. Varela, Z. Gao, N. Danné, J. P. Dupuis, B. Lounis, L. Groc, L. Cognet, Nat. Nanotechnol. 2017, 12, 238.

[47] H. Wu, R. Nißler, V. Morris, N. Herrmann, P. Hu, S.-J. Jeon, S. Kruss, J. P. Giraldo, Nano Lett. 2020, 20, 2432.

[48] J. P. Giraldo, M. P. Landry, S. Y. Kwak, R. M. Jain, M. H. Wong, N. M. Iverson, M. Ben-Naim, M. S. Strano, Small 2015, 11, 3973.

[49] S. Saghazadeh, C. Rinoldi, M. Schot, S. S. Kashaf, F. Sharifi, E. Jalilian, K. Nuutila, G. Giatsidis, P. Mostafalu, H. Derakhshandeh, Adv. Drug Del. Rev. 2018, 127, 138.

[50] P. Mostafalu, G. Kiaee, G. Giatsidis, A. Khalilpour, M. Nabavinia, M. R. Dokmeci, S. Sonkusale, D. P. Orgill, A. Tamayol, A. Khademhosseini, Adv. Funct. Mater. 2017, 27, 1702399.

[51] P. Mostafalu, A. Tamayol, R. Rahimi, M. Ochoa, A. Khalilpour, G. Kiaee, I. K. Yazdi, S. Bagherifard, M. R. Dokmeci, B. Ziaie, Small 2018, 14, 1703509.

[52] M. Ochoa, R. Rahimi, J. Zhou, H. Jiang, C. K. Yoon, D. Maddipatla, B. B. Narakathu, V. Jain, M. M. Oscai, T. J. Morken, Microsyst. Nanoeng. 2020, 6, 1.

[53] A. Memic, T. Abudula, H. S. Mohammed, K. Joshi Navare, T. Colombani, S. A. Bencherif, ACS Appl. Bio Mater. 2019, 2, 952.

[54] J. Xue, T. Wu, Y. Dai, Y. Xia, Chem. Rev. 2019, 119, 5298.

[55] E. Schnell, K. Klinkhammer, S. Balzer, G. Brook, D. Klee, P. Dalton, J. Mey, Biomaterials 2007, 28, 3012.

[56] A. F. De Faria, F. Perreault, E. Shaulsky, L. H. Arias Chavez, M. Elimelech, ACS Appl. Mater. Interfaces 2015, 7, 12751.

[57] E. Gianino, C. Miller, J. Gilmore, Bioengineering 2018, 5, 51.

[58] T. Xu, J. M. Miszuk, Y. Zhao, H. Sun, H. Fong, Adv. Healthcare Mater. 2015, 4, 2238.

[59] A. Martins, E. D. Pinho, S. Faria, I. Pashkuleva, A. P. Marques, R. L. Reis, N. M. Neves, small 2009, 5, 1195.

[60] J. Yoon, H. S. Yang, B. S. Lee, W. R. Yu, Adv. Mater. 2018, 30, 1704765.

[61] C. G. Salzmann, B. T. Chu, G. Tobias, S. A. Llewellyn, M. L. Green, Carbon 2007, 45, 907.

[62] M. S. Dresselhaus, G. Dresselhaus, R. Saito, A. Jorio, Phys. Rep. 2005, 409, 47.

[63] M. S. Dresselhaus, A. Jorio, M. Hofmann, G. Dresselhaus, R. Saito, Nano Lett. 2010, 10, 751.

[64] C. Fantini, A. Jorio, M. Souza, M. Strano, M. Dresselhaus, M. Pimenta, Phys. Rev. Lett. 2004, 93, 147406.

[65] T. Kanungo, D. M. Mount, N. S. Netanyahu, C. D. Piatko, R. Silverman, A. Y. Wu, IEEE Trans. Pattern Anal. Mach. Intell 2002, 24, 881.

[66] A. Jorio, M. Pimenta, A. Souza Filho, R. Saito, G. Dresselhaus, M. Dresselhaus, New J. Phys. 2003, 5, 139.

[67] M. J. O'Connell, E. E. Eibergen, S. K. Doorn, Nat. Mater. 2005, 4, 412.

[68] D. P. Salem, X. Gong, A. T. Liu, V. B. Koman, J. Dong, M. S. Strano, J. Am. Chem. Soc. 2017, 139, 16791.

[69] A. McLister, J. McHugh, J. Cundell, J. Davis, Adv. Mater. 2016, 28, 5732.

[70] D. Roxbury, P. V. Jena, R. M. Williams, B. Enyedi, P. Niethammer, S. Marcet, M. Verhaegen, S. Blais-Ouellette, D. A. Heller, Sci. Rep. 2015, 5, 1.

