## Supporting Information for "A Wearable Optical Microfibrous Biomaterial with Encapsulated Nanosensors Enables Wireless Monitoring of Oxidative Stress"

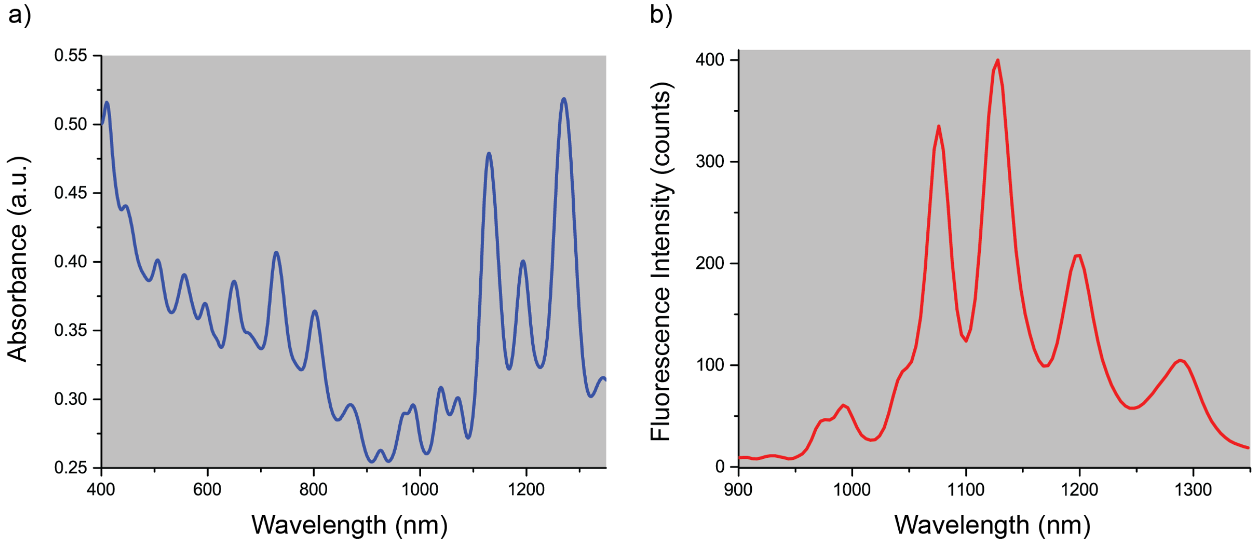


**Figure S1.** (a) Absorbance spectra, and (b) Fluorescence spectra of the $(GT)_{15}$-SWCNTs dispersion at 730 nm excitation.

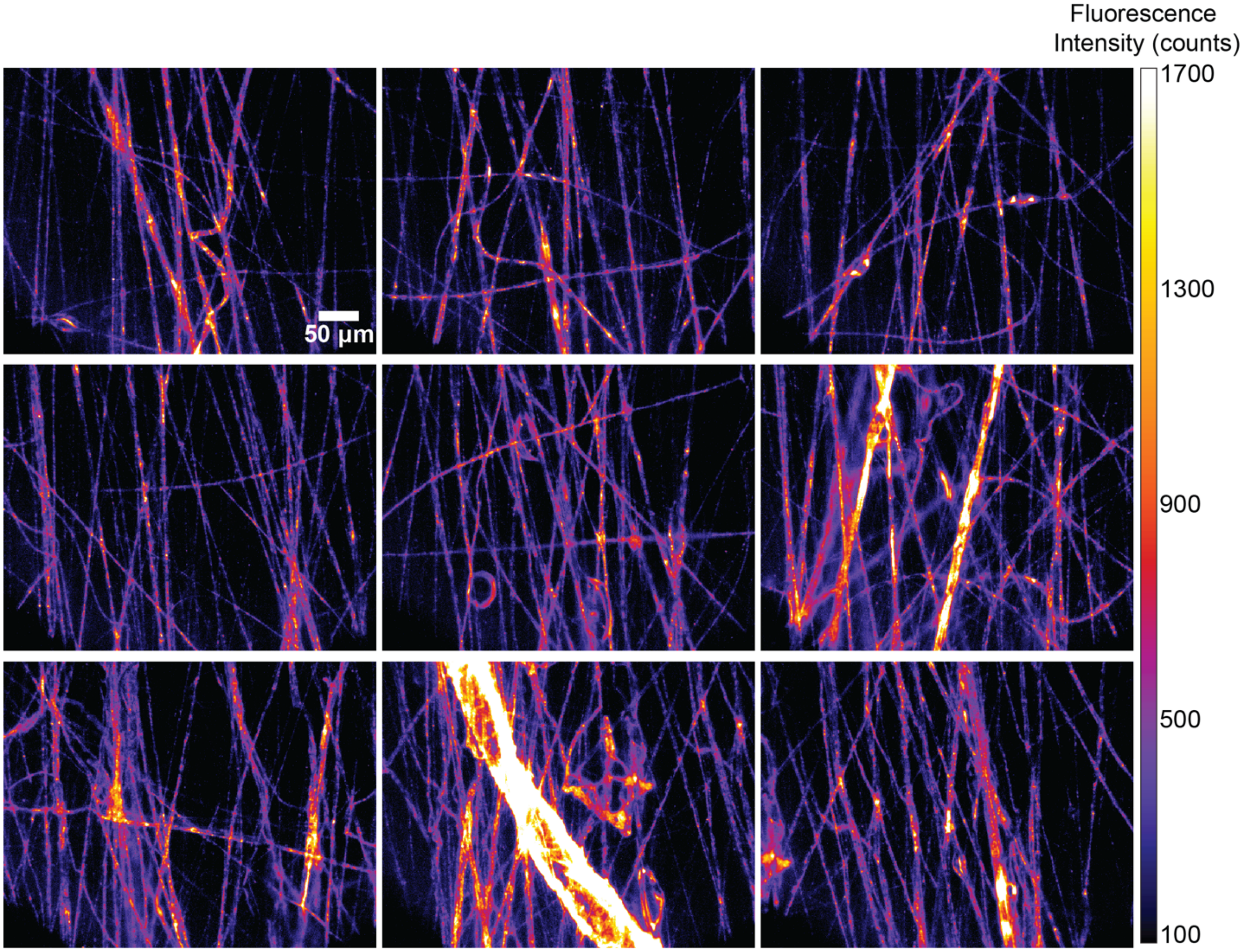


**Figure S2.** NIR broadband fluorescence images of the fibers produced with the applied voltage of 12 kV acquired at different regions.

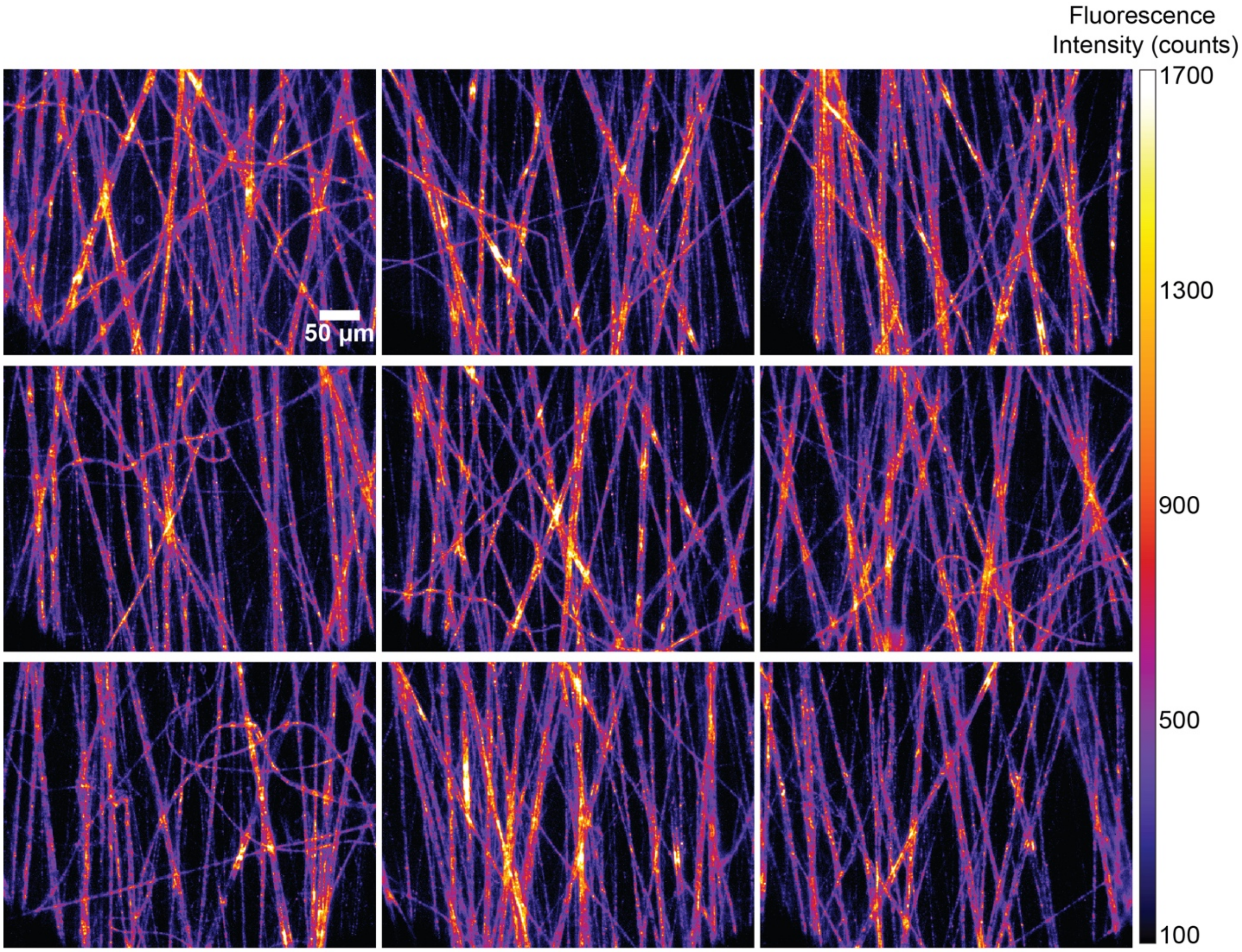


**Figure S3.** NIR broadband fluorescence images of the fibers produced with the applied voltage of 14 kV acquired at different regions.

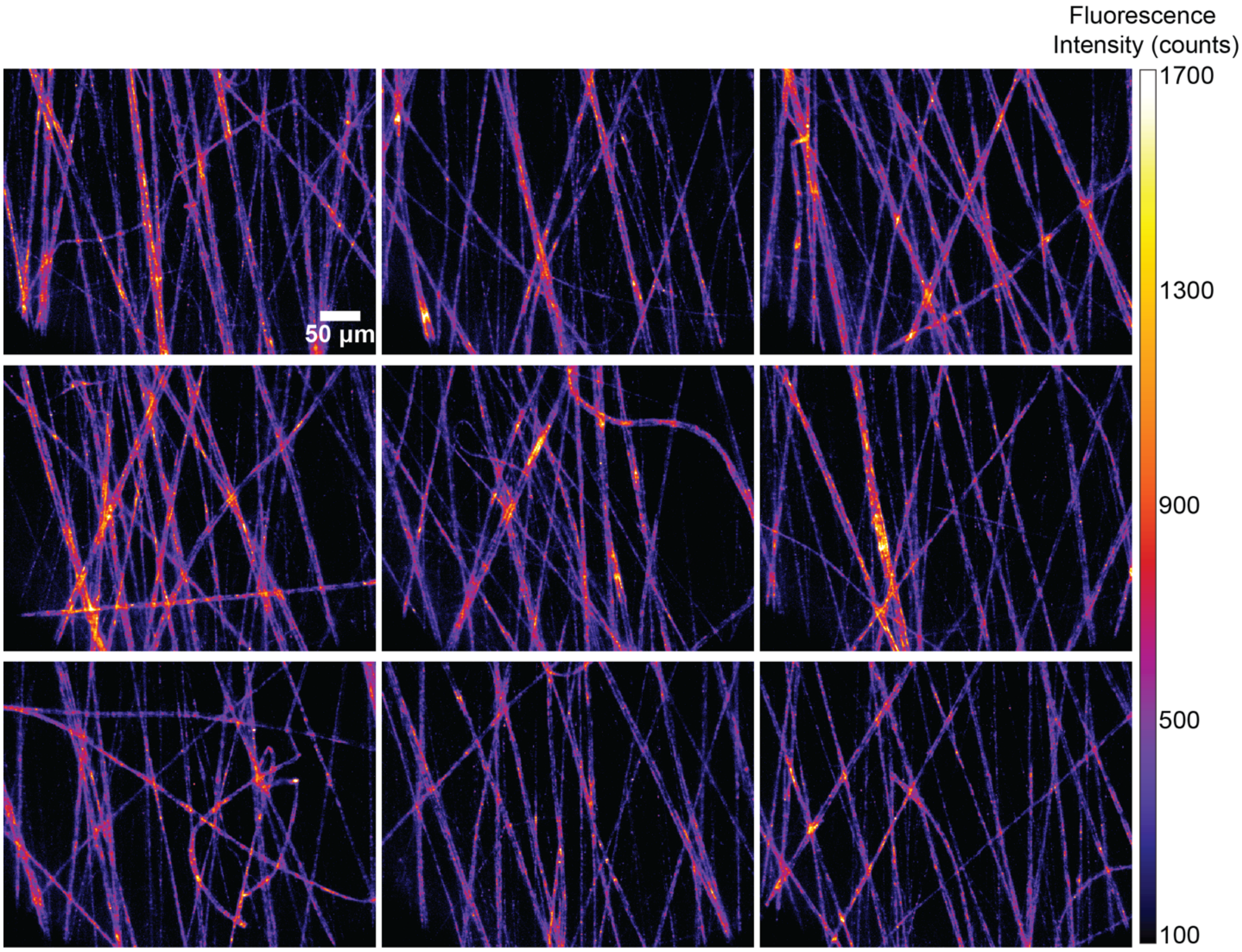


**Figure S4.** NIR broadband fluorescence images of the fibers produced with the applied voltage of 16 kV acquired at different regions.

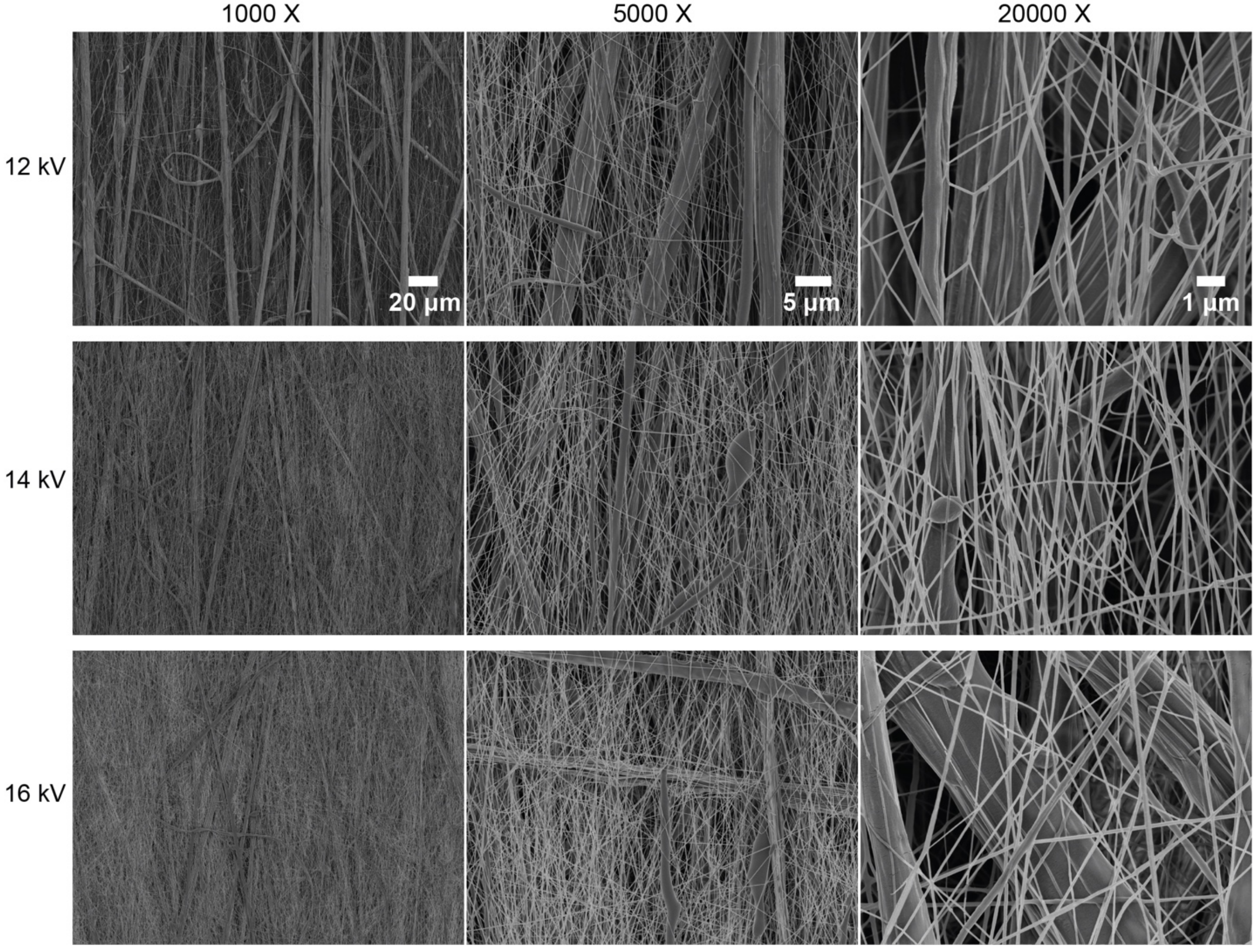


**Figure S5.** SEM images of the micro- and nanofibers fabricated with three different voltages of 12, 14 or 16 kV.

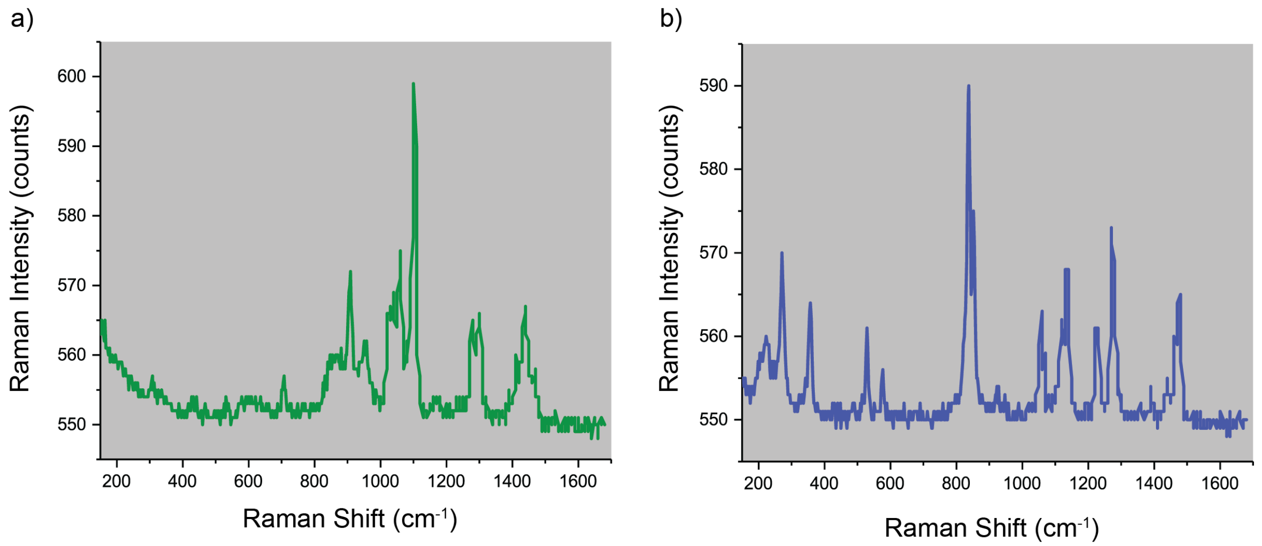


**Figure S6.** Raman spectra of (a) polycaprolactone (PCL) polymer and (b) poly(ethylene oxide) (PEO) polymer.

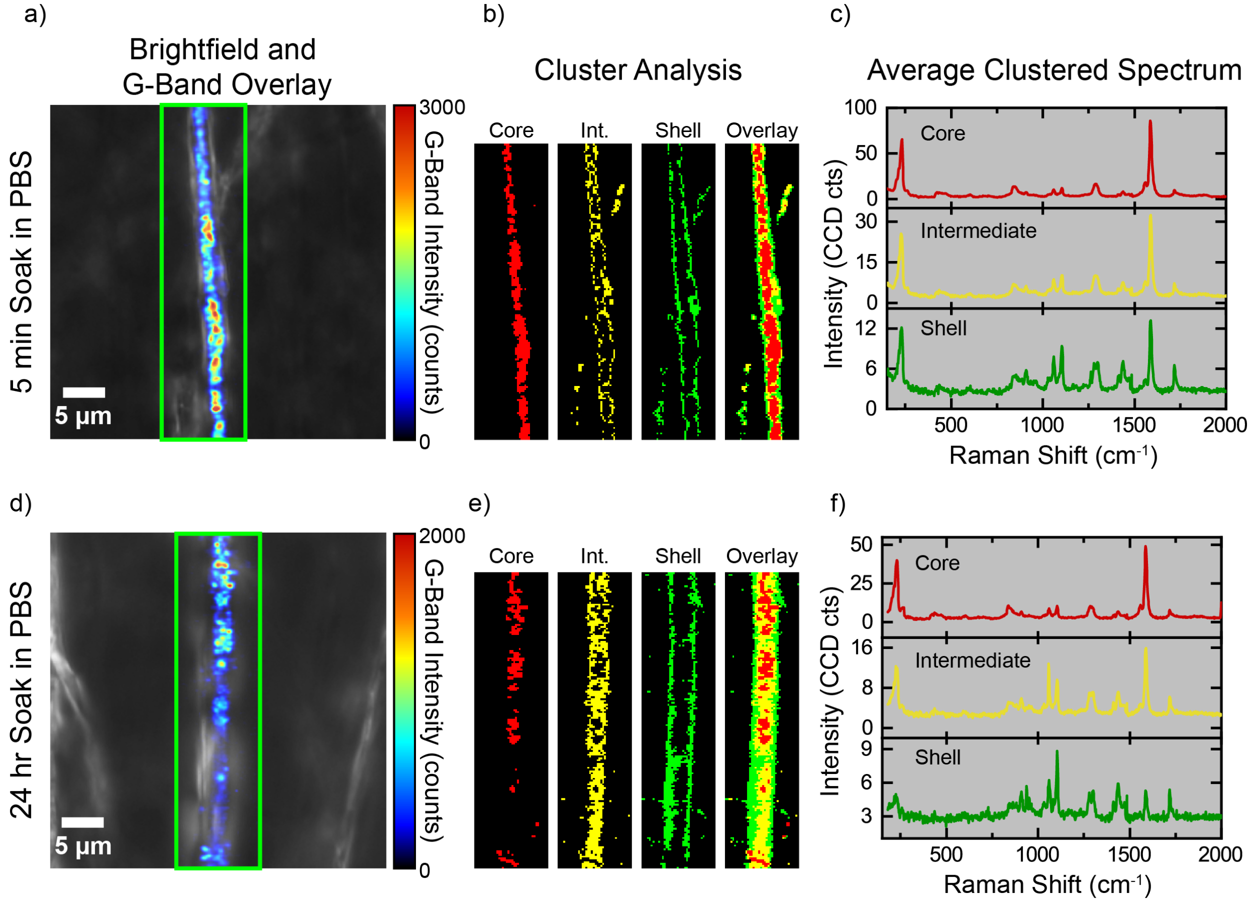


**Figure S7.** Confocal Raman microscopy of fibers soaked in PBS for 5 minutes and 24 hours. (a) and (d) The representative overlay of G-band intensity and brightfield images of fibers soaked in PBS for 5 minutes and 24 hours, respectively. (b) and (e) k-means clustering analyses of all spectra in each area scan, where $k = 4$ clusters (background clusters omitted from figure). (c) and (f) The average Raman spectra obtained from each cluster of (b) and (e).

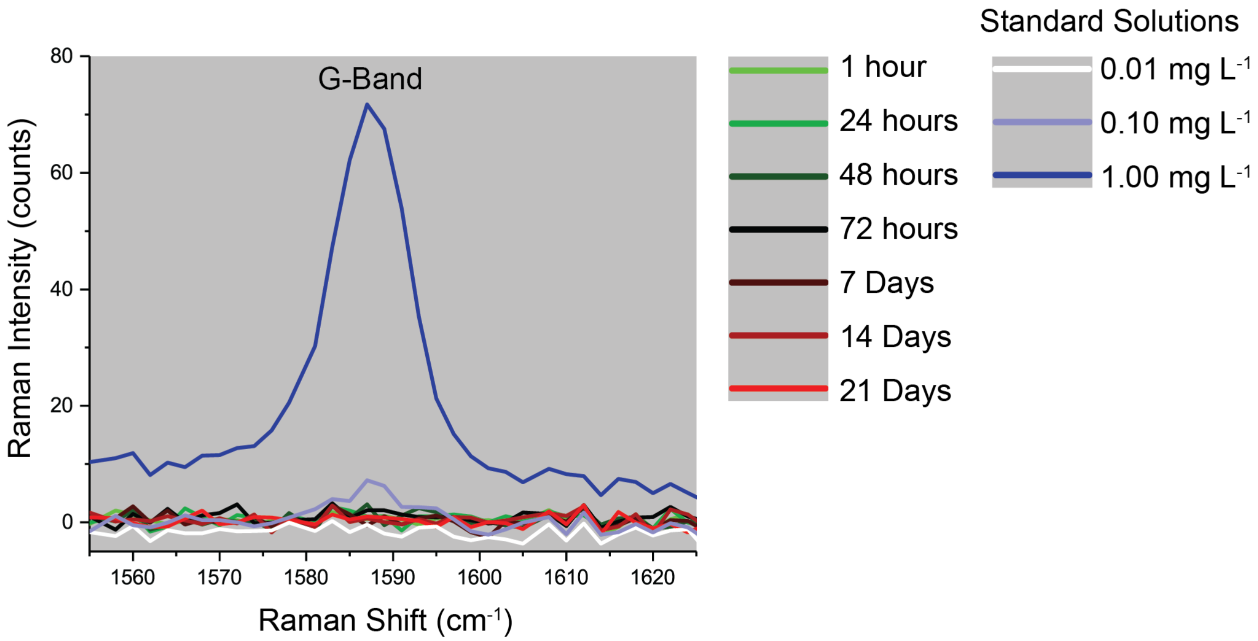


**Figure S8.** Comparing the Raman spectra of the collected PBS samples over time with that of three standard samples with known SWCNT concentrations.

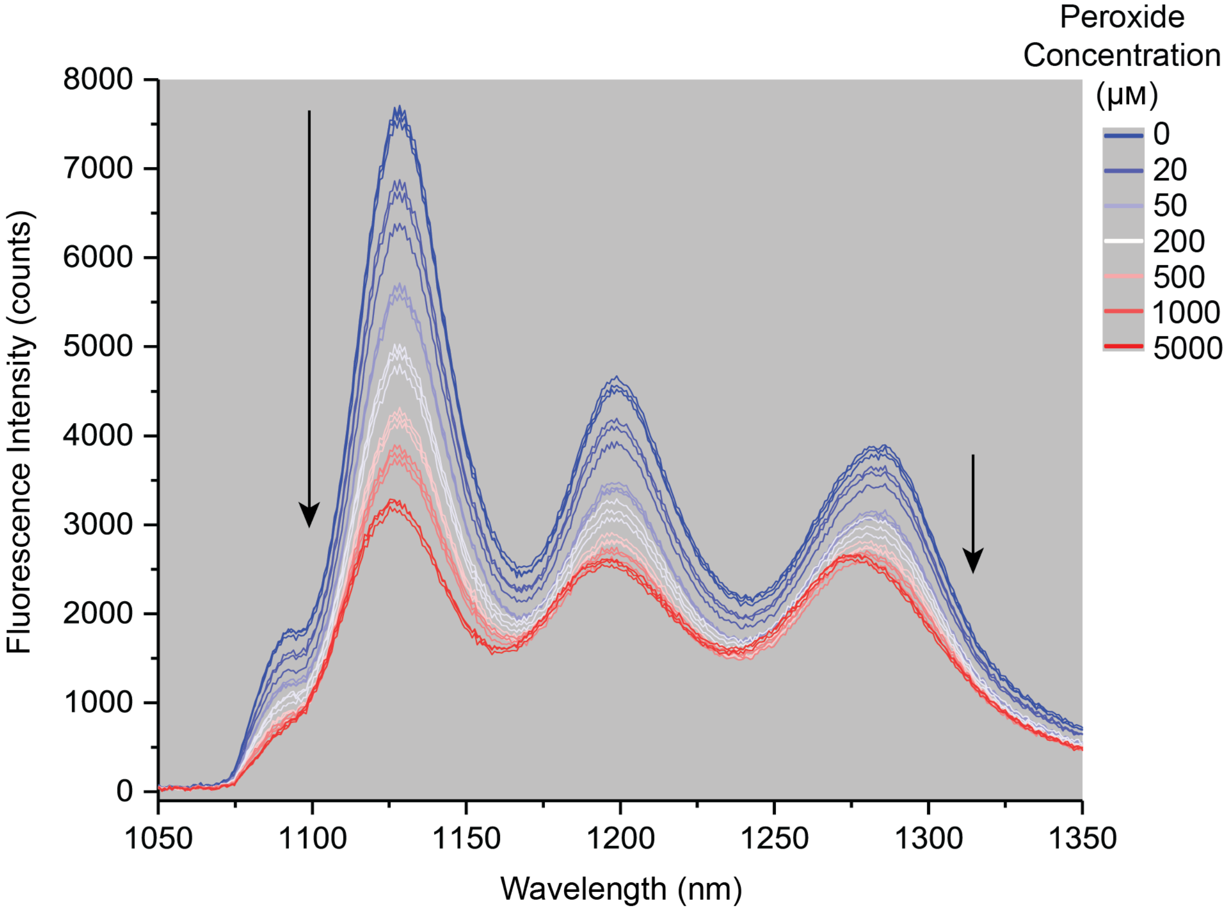


**Figure S9.** The fluorescence spectra of the microfibrous samples exposed to various peroxide concentrations. Each peroxide concentration was added to three different samples to confirm the reproducibility ($n = 3$).

**Table S1.** Linear fits to the ratiometric signal versus peroxide concentration, when plotted on a log-log scale, in the range of 5 μM-5 mM for three different time points: $R = a + bC$. Values are mean with standard error of mean (SE), from three samples for each peroxide concentration.

| Time Point | Intercept: $a \pm SE$ | Slope: $b \pm SE$ | $R^2$ |
|---|---|---|---|
| 24 hours | -0.38715 ± 0.00413 | 0.045 ± 0.00139 | 0.99241 |
| 48 hours | -0.35896 ± 0.00563 | 0.04996 ± 0.0017 | 0.99086 |
| 72 hours | -0.35609 ± 0.00508 | 0.05651 ± 0.00221 | 0.98796 |

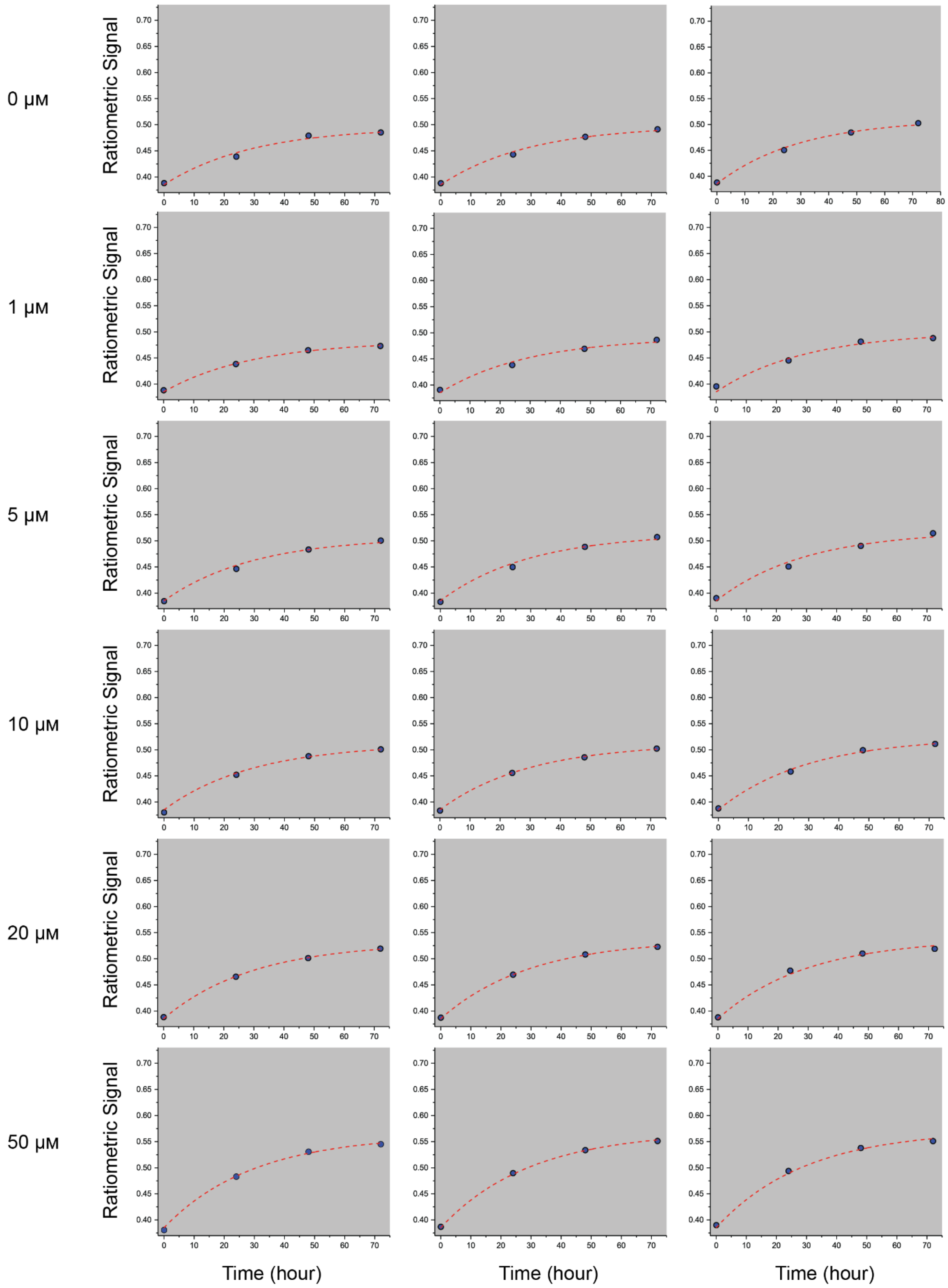
0 μM
1 μM
5 μM
10 μM
20 μM
50 μM
Ratiometric Signal
Time (hour)

**Figure S10.** Ratiometric signal as a function of time for each peroxide concentration (0-50 μM). The dashed lines indicate single exponential association fits. Each peroxide concentration was added to three different samples ($n$ = 3).

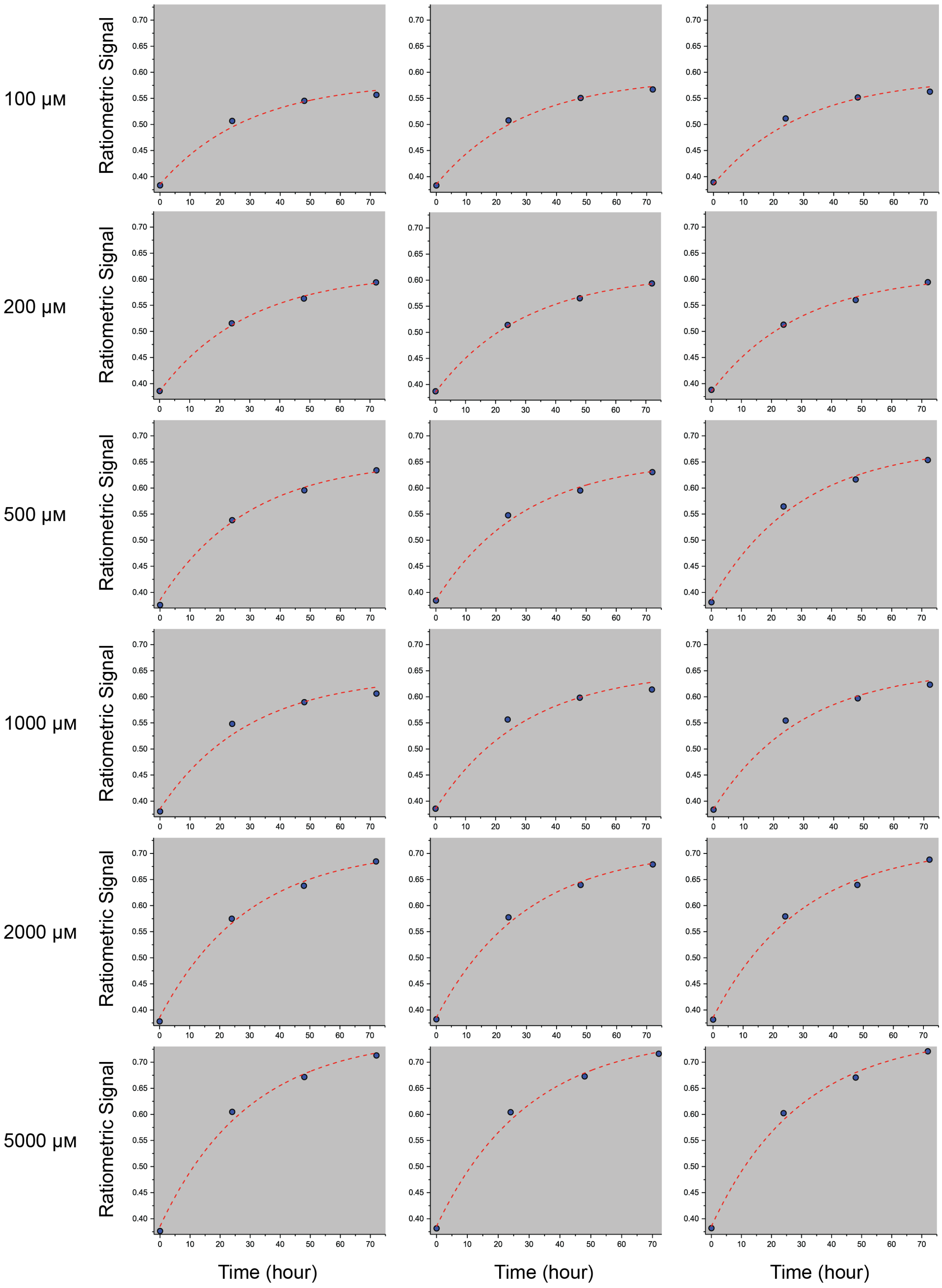

100 μM
200 μM
500 μM
1000 μM
2000 μM
5000 μM
Ratiometric Signal
Time (hour)

**Figure S11.** Ratiometric signal as a function of time for each peroxide concentration (100-5000 μM). The dashed lines indicate single exponential association fits. Each peroxide concentration was added to three different samples ($n$ = 3).

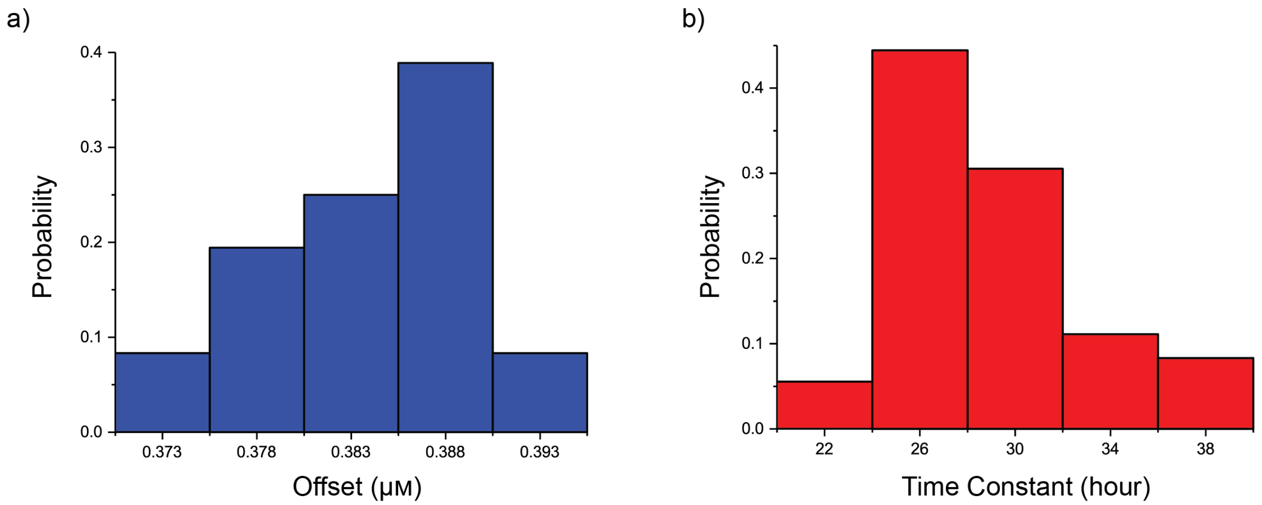

**Figure S12.** Histograms indicating the distribution of (a) Offset and (b) Time constant extracted from the single exponential association fits of the ratiometric signal versus time, for all examined concentrations of peroxide.

**Table S2.** Single exponential association fits to the ratiometric signal versus time for each peroxide concentration: $R = R_0 + A\left(1 - e^{-t/\tau}\right)$. Values are mean with standard error of mean (SE), from three samples for each peroxide concentration.

| Concentration [μM] | Offset: $R_0 \pm SE$ | Pre-Exponential Factor: $A \pm SE$ | Time Constant: $\tau$ [s] $\pm SE$ | $R^2$ |
|---|---|---|---|---|
| 0 | 0.3887 ± 1.41E-04 | 0.11629 ± 0.00454 | 33.39577 ± 0.12185 | 0.99365 |
| 1 | 0.39137 ± 0.00204 | 0.10605 ± 0.00498 | 35.87988 ± 1.9239 | 0.99477 |
| 5 | 0.38787 ± 0.00201 | 0.12837 ± 0.00342 | 32.9493 ± 1.61385 | 0.99214 |
| 10 | 0.38456 ± 0.00184 | 0.13064 ± 0.00398 | 29.39384 ± 1.29415 | 0.99826 |
| 20 | 0.38661 ± 0.00118 | 0.1504 ± 0.00228 | 30.07832 ± 1.24041 | 0.99739 |
| 50 | 0.3848 ± 0.00202 | 0.183 ± 0.00281 | 28.94404 ± 0.86832 | 0.99827 |
| 100 | 0.38052 ± 0.00158 | 0.20274 ± 0.00258 | 25.86262 ± 0.74715 | 0.99279 |
| 200 | 0.38713 ± 8.67E-04 | 0.2268 ± 7.08E-04 | 30.33552 ± 0.45051 | 0.99862 |
| 500 | 0.37924 ± 0.00139 | 0.27885 ± 0.00903 | 26.99243 ± 0.356 | 0.99615 |
| 1000 | 0.3743 ± 0.00163 | 0.26379 ± 0.00418 | 23.58728 ± 0.76148 | 0.98822 |
| 2000 | 0.37977 ± 7.78E-04 | 0.32639 ± 0.00155 | 27.53061 ± 0.21989 | 0.99676 |
| 5000 | 0.37714 ± 0.00234 | 0.36604 ± 8.48E-04 | 26.72116 ± 0.70784 | 0.99657 |
| | Avg = 0.383 ± 9.28E-4 | | Avg = 29.305 ± 0.615 | |

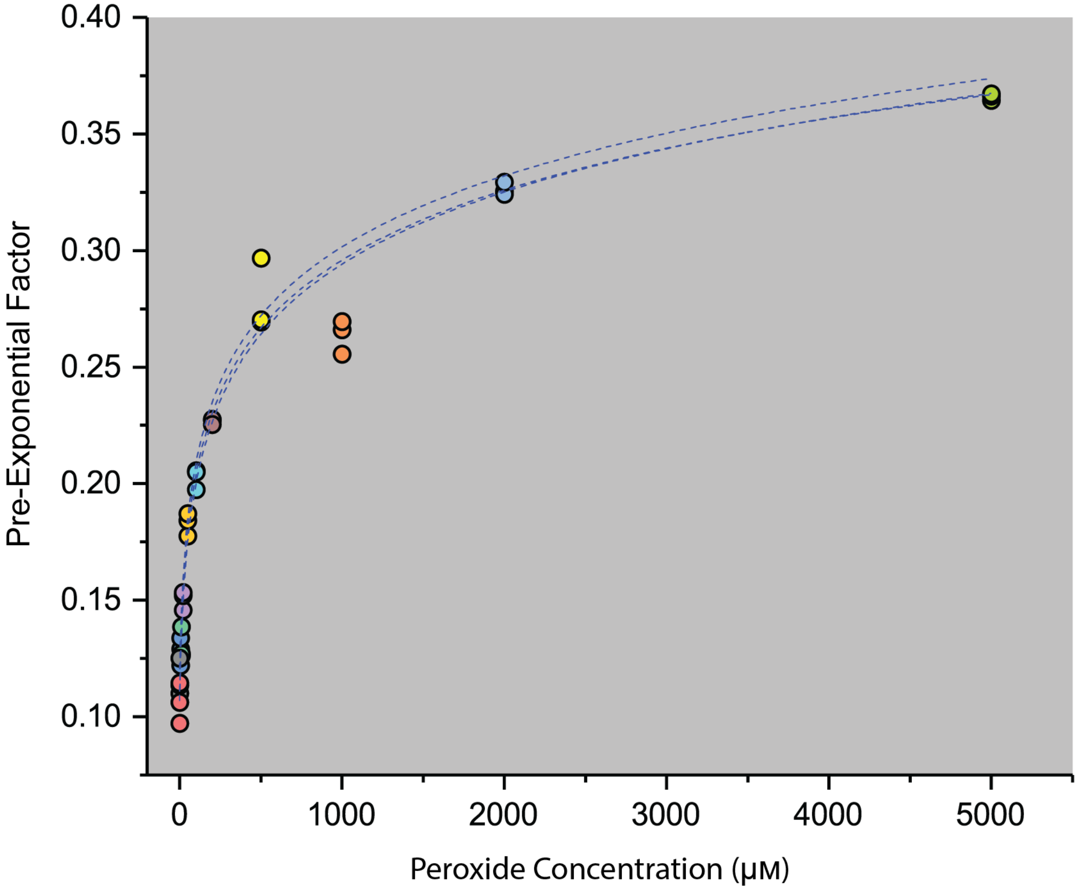


**Figure S13.** The pre-exponential factor extracted from the single exponential association fits and plotted as a function of peroxide concentration. The dashed lines indicate the power fits. Each peroxide concentration was added to three different samples ($n = 3$).

**Table S3.** Power fits to pre-exponential factor versus peroxide concentration:

$$A = (C + C_0)^p - A_0$$

| | $C_0$ [μM] | $p$ | $A_0$ | $R^2$ |
|---|---|---|---|---|
| 1 | 10.69924 | 0.03495 | 0.97934 | 0.9986 |
| 2 | 9.81501 | 0.03408 | 0.96994 | 0.99763 |
| 3 | 11.87454 | 0.03458 | 0.96881 | 0.98754 |
| Avg | 10.796 ± 0.596 | 0.035 ± 2.52E-4 | 0.972 ± 3.34E-3 | |

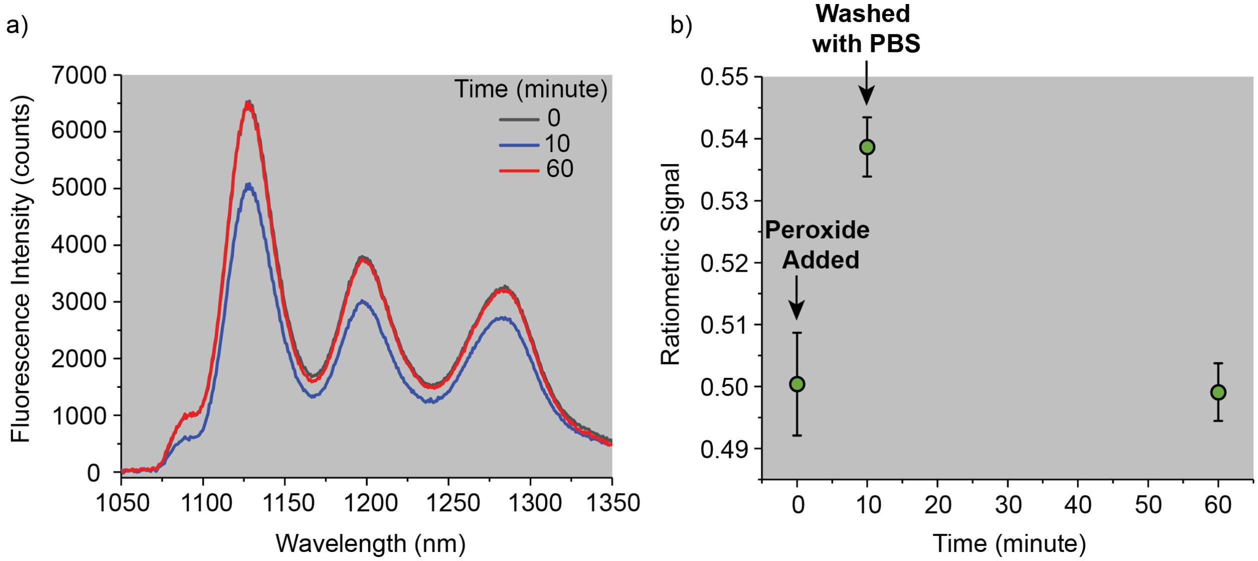


**Figure S14.** Reversible peroxide detection using the optical microfibrous textiles. (a) The NIR fluorescence spectra of the samples right before adding 200 μM peroxide (0 min), 10 minutes after peroxide addition and 50 minutes after removing the peroxide (the samples were washed with PBS after 10 minutes of exposure to peroxide). (b) The ratiometric signal at the three time points mentioned in part a. Mean values were obtained by repeating each condition three times ($n = 3$), and the error bars represent the standard deviation.

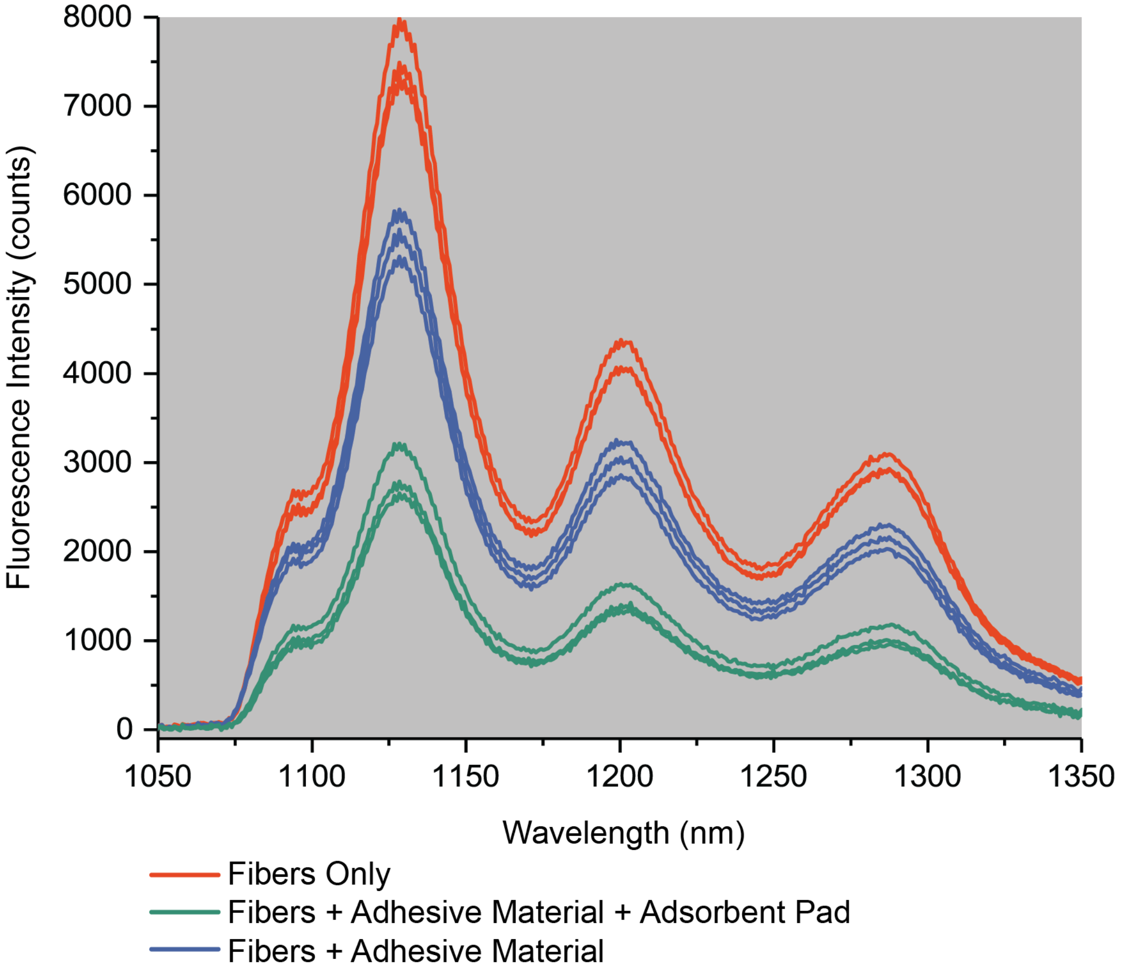


**Figure S15.** Comparison of the fluorescence spectra of microfibers alone, through adhesive bandage material, or through both adhesive material plus an adsorbent pad (complete bandage). The fluorescent spectra were acquired from three different samples per each condition ($n$ = 3).
